# *Talaromyces marneffei* suppresses human macrophages inflammatory by producing the truncated protein NCOR2-013 via TUT1-regulated alternative splicing

**DOI:** 10.1101/2022.07.11.499655

**Authors:** Wudi Wei, Gang Wang, Hong Zhang, Xiuli Bao, Sanqi An, Qiang Luo, Jinhao He, Lixiang Chen, Yuxuan Liu, Chuanyi Ning, Jingzhen Lai, Zongxiang Yuan, Rongfeng Chen, Junjun Jiang, Li Ye, Hao Liang

**Affiliations:** Guangxi Key Laboratory of AIDS Prevention and Treatment, School of Public Health, Guangxi Medical University, Nanning 530021, Guangxi, China; Guangxi-ASEAN Collaborative Innovation Center for Major Disease Prevention and Treatment, Life Sciences Institute, Guangxi Medical University, Nanning 530021, Guangxi, China; Nursing College, Guangxi Medical University, Nanning 530021, Guangxi, China; Guangxi Biobank, Life Sciences Institute, Guangxi Medical University, Nanning 530021, Guangxi, China

**Keywords:** *Talaromyces marneffei*, macrophages, NCOR2-013, TUT1, alternative splicing

## Abstract

*Talaromyces marneffei* (*T. marneffei*) immune-escaping is an important factor for high mortality of talaromycosis. It is currently known that *T. marneffei* performs these functions through a variety of strategies, however, the role of alternative splicing (AS) in this process is poorly understood. Here we depicted the AS landscape in the macrophage upon *T. marneffei* infection via high-throughput RNA sequencing. Moreover, we identified a truncated protein of NCOR2/SMRT, namedly NCOR2-013, was significantly upregulated upon *T. marneffei* infection. Mechanistic analysis indicates that NCOR2-013 forms a co-repression complex with TBL1XR1/TBLR1 and HDAC3, thereby inhibiting JunB-mediated transcriptional activation of pro-inflammatory cytokines via the inhibition of histone acetylation. Also, we identified TUT1 as the AS regulator that involved in facilitating *T. marneffei* immune evasion via regulation of NCOR2-013 production. Collectively, the findings indicate that *T. marneffei* escapes macrophages killing through the TUT1-mediated the alternative splicing of NCOR2-013, which providing a new insight into the molecular mechanisms of *T. marneffei* immune evasion, and a potential targets for talaromycosis therapy.

## INTRODUCTION

Talaromycosis (formerly penicilliosis) is caused by the thermally dimorphic fungus *Talaromyces marneffei* (*T*.*marneffei*) that is one of the important causes of death among AIDS patients in epidemic area. Talaromycosis is a regional high-incidence opportunistic infection that is endemic throughout Southern China, Southeast Asia, and Northeastern India (1,2). The number of talaromycosis cases has rapidly increased due to the HIV epidemic, ranking 3^rd^ as the most common HIV-associated opportunistic infections and accounting for up to 16% of HIV admissions and is a leading cause of death in patients with advanced HIV disease in Thailand, Vietnam, and southern China (1–3). 288,000 cases have been reported in 33 countries to the end of 2018, with an estimated 17,300 cases (95% CI 9,900-23,700) and 4,900 deaths (2,500-7,300) each year(4).Therefore, talaromycosis has become a serious local public health problem, which is proposed to be included in the “List of Priority Fungal Pathogens” by WHO and is called to be recognised as a neglected tropical infectious disease (5,6).

*T*.*marneffei* immune evasion is an important reason for the poor prognosis of talaromycosis, previous studies have shown that *T*.*marneffei* can survive in host macrophages. Felix et al found that zebrafish embryos macrophages protect *T*.*marneffei* from neutrophil fungicidal activity during infection (7). Thus, it is obvious that *T*.*marneffei* may have a set of macrophage-based immune evasion strategies in the human body. As a facultative intracellular fungus, *T*.*marneffei* are readily phagocytosed by resident macrophages after invasion (1,8). Paradoxically, as the body’s first line of defense against pathogen invasion, macrophages provide *T*.*marneffei* with a niche to evade immune killing, which known as the “macrophage paradox” (9). In fact, several pathogens have similar strategy, such as Mycobacterium tuberculosis, Candida albicans, etc (9–11). Therefore, to understand the molecular mechanism of *T*.*marneffei* immune evasion is of great significance for the treatment of talaromycosis patients and the reduction of mortality. Previous studies have demonstrated that *T. marneffei* may escape macrophage killing to proliferate inside macrophage by inducing M2-like polarization via regulating SOCS3-STAT6, TLR9, Jun1/2, and p38 signaling pathways, also the LncRNA (12–15). However, a deeper understanding of the mechanisms are still urgently needed.

Alternative splicing (AS) is a ubiquitous mechanism for regulating gene expression that allows a single gene to produce multiple unique mRNAs. Previously, it was reported that approximately 90%–95% of human genes undergo several level of AS, almost 37% of which were proved to generate multiple protein isoforms that may play distinct biological functions (16–19). Generally, AS includes five categories of events, namely intron retention (RI), exon skipping (ES), 5 SS alternative splicing (A5SS), 3 SS alternative splicing (A3SS), and mutually exclusive exon splicing (MXE), generates multiple mature mRNA isoforms from the same pre-mRNA (20). Thus, AS plays an important role in the complexity and functional diversity of the proteome. Bacterial pathogens are known to strategically modulate specific host factors via regulating AS to facilitate its own replication in host cells (21–23). For instance, Mtb can survive by limiting the phagosome maturationthe via producting abundance of truncated RAB8B variant (22). As a highly dynamic and complex process, AS is mainly regulated by a broad array of RNA binding proteins (RBP). Mechanistically, RBP binds to target pre-mRNA through specific motifs to mediate the occurrence of AS process (16,18). Intriguingly, the global changes in gene AS events upon infection of macrophages with *T*.*marneffei* are not extensively understood, and there is no clear description of whether specific spliceosome components could help *T*.*marneffei* immune evasion.

In this study, we performed high-throughput RNA sequencing and depicted global view of the changes in gene AS events upon *T*.*marneffei* infection. Moreover, we identified a TUT1-modulated AS event of NCOR2-013 involved in facilitating *T*.*marneffei* immune evasion. In summary, our data show that the alternative splicing of NCOR2-013 promotes *T*.*marneffei* immune evasion regulating by TUT1, implying that NCOR2-013 has potential as a therapeutic target for talaromycosis therapy.

## MATERIALS AND METHODS

### Reagents and antibodies

Human IFN-γ and,GM-CSF recombinant protein was obtained from Sinobio Biotechnology, and the LPS from E. coli was from Sigma-Aldrich. The IRDye 680RD Donkey anti-Mouse and IRDye 800CW Donkey anti-Rabbit antibody was purchased from LI-COR Biosciences. Anti-rabbit IgG (H+L), F(ab’)2 Fragment (Alexa Fluor® 647 Conjugate), Anti-FLAG (D6W5B), anti-PCNA (D3H8P), anti-β-Actin (8H10D10), anti-Normal Rabbit IgG, anti-HDAC3 (D2O1K), anti-TBL1XR1/TBLR1 (D4J9C), anti-JunB (C37F9), anti-Phospho-JunB (Thr102/Thr104) (D3C6), anti-JunD (D17G2), anti-c-Jun (60A8), anti-Phospho-c-Jun (Ser73), anti-Acetyl-Histone H3 (Lys27) (D5E4) and anti-GAPDH (D16H11) were were purchased from Cell Signaling Technology. Anti-NCOR2 was obtained from Novus Biologicals. Anti-TUT1 (PA5-50151) was obtained from Invitrogen.

### Fungal strain

*T. marneffei* strain (ATCC18224) was purchase from American Type Culture Collection (ATCC) and used for all experiments. *T. marneffei* was firstly grown in potato dextrose agar medium (Sigma, USA) and the conidia were obtained after 7-10 days of incubation at 27°C. Spores were filtered through sterile glass wool (Corning, USA), and the filtrate was centrifuged to produce a spore pellet. Conidia were resuspended at 10^7^ condidia/ml in sterile PBS for the subsequent experiments.

### THP-1 macrophages and human monocyte-derived macrophage (hMDM)

THP-1 cells were purchase from American Type Culture Collection (ATCC) and cultured in RPMI 1640 medium (Solarbio, China) containing 10% fetal bovine serum (FBS, Gibco, USA) and 1% penicillin/streptomycin (Solarbio, China). Moreover, THP-1 cells were incubated with 100 ng/ml phorbol myristate acetate (PMA, Sigma, USA) for 48h to differentiate into macrophages. Human monocyte-derived macrophage (hMDM) were obtained from human peripheral blood mononuclear cells (PBMCs) of healthy adult donor. PBMCs were isolated by density gradient centrifugation (Ficoll 1.077 g/mL, Sigma, USA), diluted at 10^6^ cells/mL in Dulbecco’s Modified Eagle’s Medium (DMEM, Solarbio, China) supplemented with 10% FBS and 1% penicillin/streptomycin. In order to obtain fully differentiated human primary macrophages, the PBMCs were cultured for 7 days in presence of 25ng/mL recombinant human macrophage colony-stimulating factor (rh-MCSF, Sino Biological, China). Cells were cultured at 37°C in a humidified atmosphere with 5% CO_2_.

### Lentivitus transfection

Prior to transfection, THP-1 cells were seeded in 96-well plates in an incubator at 37°C overnight, and human derived macrophages were differentiated from PBMCs in 96-well plates. Then, siRNA-TUT1 lentivirus, Flag-TUT1-overexpressing lentivirus, Flag-NCOR2-013-overexpression lentivirus and their negative control (NC) lentivirus were added into the plates, and cultured in serum-free RPMI 1640 (for THP-1 cells) and DMEM (for THP-1 macrophages and hMDMs) media. The lentivirus volume equals (MOI × number of cells) / lentivirus titer. After 48h of transfection, the medium containing lentivirus was removed and the fresh complete medium was added.

### Primers designing and quantitative real-time PCR (qRT-PCR) analysis

Primers were designed using primer-BLAST and and Primer Premier 5 software to amplify the exon regions of NCOR2, IL-1β, TNF-ɑ, etc. Note that all these primers were designed to amplify exon region. Primers were optimized prior to quantification experiments using qPCR. The sequences were shown in Supplementary Appendix. qRT-PCR was performed using SYBR Premix ExTaq™ (TAKARA, Japan) by StepOnePlus Real-time PCR system (Thermo Fisher Scientific, USA). Relative gene expression was defined as a ratio of target gene expression versus GAPDH genes expression. Data were analyzed with 2^−ΔΔ Ct^ method. The primer design and reactions for ChIP-qPCR were the same as above.

### Western blot

The nuclear/cytoplasmic and total proteins in THP-1-derived macrophages were extracted, respectively, using the nuclear protein and cytoplasmic protein extraction kit (#P0028, Beyotime, China) and cell lysis buffer (#9803, CST, USA) according to the manufacture’s instruction. The protein concentration was detected using a BCA protein assay kit (#P0012S, Beyotime, China). Protein expression levels were measured by western blot. Equal amount of protein were separated with 10-12% SDS-PAGE (BOSTER, China) and transferred onto a PVDF membrane (Bio-Rad, USA). After blocking with 5% nonfat milk (BD Biosciences, USA) for 2h, the membrane was incubated with primary antibody overnight with mild shake at 4 °C. After that, the membrane was washed with TBST three times and incubated with the corresponding secondary antibody for 1h. Images were performed using the Odyssey CLX two-color infrared laser imaging system (Odyssey LI-COR, USA).

### Colony-forming units (CFU)

Colony forming units is a possible method to quantify the amount of *T. marneffei* living in macrophages. The supernatant was discarded and the macrophages were washed twice with PBS, then the intracellular *T. marneffei* was collected by lysing the cells with sterile water, the volume was made up to 200µl. Afterwards, the fungal suspension was diluted with sterile water into 3 concentration gradients (10^−1^, 10^−2^, and 10^−3^), and inoculated (5µl) on yeast extract peptone dextrose solid medium, respectively. The plates were incubated at 37°C for 24-48 h. The rationale for various concentration gradients is to obtain the number of colony between 30 and 300. If the counts were smaller/larger than these range, then the counts from another dilution gradients were used. The fungal load of macrophages was calculated with the following formula: Fungal load = (number of colonies × dilution gradient) × 40.

### Immunofluorescent staining

Immunofluorescence was performed to detect the NCOR2-013 subcellular localization. THP-1 macrophages were plated onto cover slips. The cells were fixed with 4% paraformaldehyde solution for 20 min, and then washed 3 times with PBS. For permeabilization, cells were incubated with 1% Triton X-100 PBS solution for 20min at room temperature, and again washed 3 times with PBS as method mentioned above. After blocking with 5% nonfat milk for 1h, the samples were incubated overnight at 4°C with the desired primary antibody (1:200), washed with PBS, and then incubated with a specific fluorescence-conjugated secondary antibody for 1h in a light-protected chamber at room temperature. Then, DAPI (4,6-diamidino-2-phenylindole) was used to label nuclei. Immunofluorescence signals were detected using fluorescence microscopy (Nikon, Japan).

### Cytometric bead array (CBA) assay

Cytokine levels were analyzed by cytometric bead array. After the NCOR2-013-overexpression macrophages and NC vector were co-cultured with *T. marneffei* for 24h, the supernatants were collected for detection of secreted cytokines. TNF-ɑ and IL-1β cytometric bead array assays were performed according to the manufacturers instructions (BD Biosciences, USA). Briefly, 50µl supernatants were mixed with 50µl capturing beads and 50µl detection reagents of the kit, then incubated for 2h at room temperature in the dark. Samples were washed and re-suspended in 200 µl wash buffer and analyzed by flow cytometry (Beckman Coulter, USA).

### Immunoprecipitation (IP) and Co-Immunoprecipitation (Co-IP)

Cell lysates were prepared in RIPA (for immunoprecipitation) or Co-IP buffer (for co-immunoprecipitation) supplemented with phosphatase and protease inhibitors. Lysates were pre-cleared with either anti-mouse or anti-rabbit IgG agarose beads for 30min. Proteins were precipitated with 5µg of primary antibodies for 1h and subsequently collected with anti-mouse (or anti-rabbit) IgG agarose beads by overnight incubation. Bead-protein complex was washed 6 times with PBS and immunoprecipitates were subsequently eluted and separated by SDS-PAGE gel electrophoresis, followed by Western blotting.

### Chromatin immunoprecipitation (ChIP)

Chromatin immunoprecipitation assays were carried out using the Agarose ChIP kit from Thermo Scientific, according to the manufacturer’s guidelines. For immunoprecipitation, 5µg of each antibody or negative control IgG were used. qRT-PCR was performed on the purified DNA and input DNA, using SYBR Premix ExTaq™ by StepOnePlus Real-time PCR system. The sequences were shown in Supplementary Appendix. Data were analyzed with 2 ^−ΔΔ Ct^ method. Each PCR was repeated at least twice, and the mean value of technical replicates was recorded for each biological replicate. Data from three independent experiments were collected, and error bars represent the standard deviation (SD) from independent experiments.

### RNA-binding protein Immunoprecipitation (RIP)

RIP assay and RNA pull down assay were performed using RIP RNA-Binding Protein Immunoprecipitation kit (Sigma, USA) and Magnetic RNA-Protein Pull-Down Kit (Thermo Fisher Scientific, USA) according to the manufacturer’s protocol, respectively. The primer design and reactions were the same as above.

### Liquid chromatography-mass spectrometry (LC-MS)

We used DE-MS to screen for proteins that bound to NCOR2-013. Briefly, NCOR2-013/Flag and isotype control IgG antibodies were used to immunoprecipitate proteins, respectively, and NCOR2-013 protein interactors were identified by in-gel digestion and LC-MS analysis. The LC-MS analysis was performed by GeneChem Co., Ltd (Shanghai, China).

### RNA-Seq and function enrichment analysis

The library preparation and RNA-Seq was performed by GeneChem Co., Ltd (Shanghai, China). Briefly, a total of 200ng of RNA in a 50μl volume was used for preparation of paired-end sequencing libraries. Afterward, samples were subjected to quality control, cluster generation and sequencing. The reads were de-multiplexed and converted to FASTA format. Finally, alignment and QC were performed. Transcript counts were normalized using conditional quantile normalization for transcript length and GC content. We then tested for differential expression between the macrophages and T. marneffei-infected macrophages using a two-sided Wald test in DESeq2. The statistical significance was set to |log2 fold of change| > 0.4 and adjusted *P* value < 0.05. The heatmap was constructed using the pheatmap R package to inflect the expression intensity and direction of the DEGs. The potential function of DEGs among different times infected with T. marneffei, modules enrichment analyses were predicted using Clusterprofiler R package. Genes with |log2 fold of change| > 0.4 and adjusted P value < 0.05 were selected for Kyoto Encyclopedia of Genes and Genomes (KEGG) and Gene Ontology (GO) analysis. Gene set enrichment analysis (GSEA) was performed using the ‘clusterProfiler’ R package.

### Statistical analysis

Each experiment was repeated at least 3 times and performed separately. Comparisons were performed by Student’s t test or one-way ANOVA analysis, and results were presented as the mean ± SD. *P* values less than 0.05 were considered statistically significant. Normality of data was tested using the Shapiro-Wilk test. For RNA-seq analysis, DEGs were calculated in DESeq2 using the Wald test with Benjamini-Hochberg correction to determine FDR. All analyses were performed using SPSS 23.0, Graphpad Prism 8.0 and R studio.

### Ethics statement

The studies involving human participants were reviewed and approved by Ethics and Human Subjects Committee of Guangxi Medical University (Ethical Review No. 20220018). The patients/participants provided their written informed consent to participate in this study.

## RESULTS

### Profiles of alternative splicing in *T. marneffei*-infected THP-1 macrophages

To investigate the potential function of alternative splicing (AS) events involved in *T. marneffei*-infected macrophages, we performed RNA sequencing (RNA-seq) in *T. marneffei*-infected or non-infected THP-1 macrophages. Collectively, 75,939 AS events were found in *T. marneffei*-infected macrophages, of which 1757 were statistically significant. The proportion of ES in significant AS events was 60.33%, which was the highest compared with other types of AS events (Fig. 1A). The UpSet plot showed that the largest group was contained in ES type (763), followed by RI (166), and only 6 genes had 3 AS types. Most of the genes were involved in one type of AS event (Fig. 1B). Afterward, the DEGs obtained from RNA-seq analysis were matched with 874 AS events to discover potential AS events associated with biological functions after *T. marneffei* infection. The result showed that ES-type AS events occurred in 27 DEGs, with 5, 3, 7, and 2 for RI, A3SS, A5SS, and MXE types, respectively (Fig. 1C). KEGG enrichment analysis of ES-type AS genes showed significantly enrichment of several pathogen immunity-related pathways, with the top-ranked pathway highlighting Epstein-Barr virus infection, influenza A and measles (Fig. S1). GO and KEGG pathway functional enrichment analyses of other AS event types were showen in Supplementary Figure 2. Finally, we selected ES-type AS events with FDR < 0.05 and |Lnclevel| > 0.5 for further enrichment analysis, and 6 genes were found in the main pathways (NCOR2, NR1H3, PPARA, IRF3, MAP2K3 and TLR6) (Fig. 1D, Fig. S3A).

**Figure 1.**
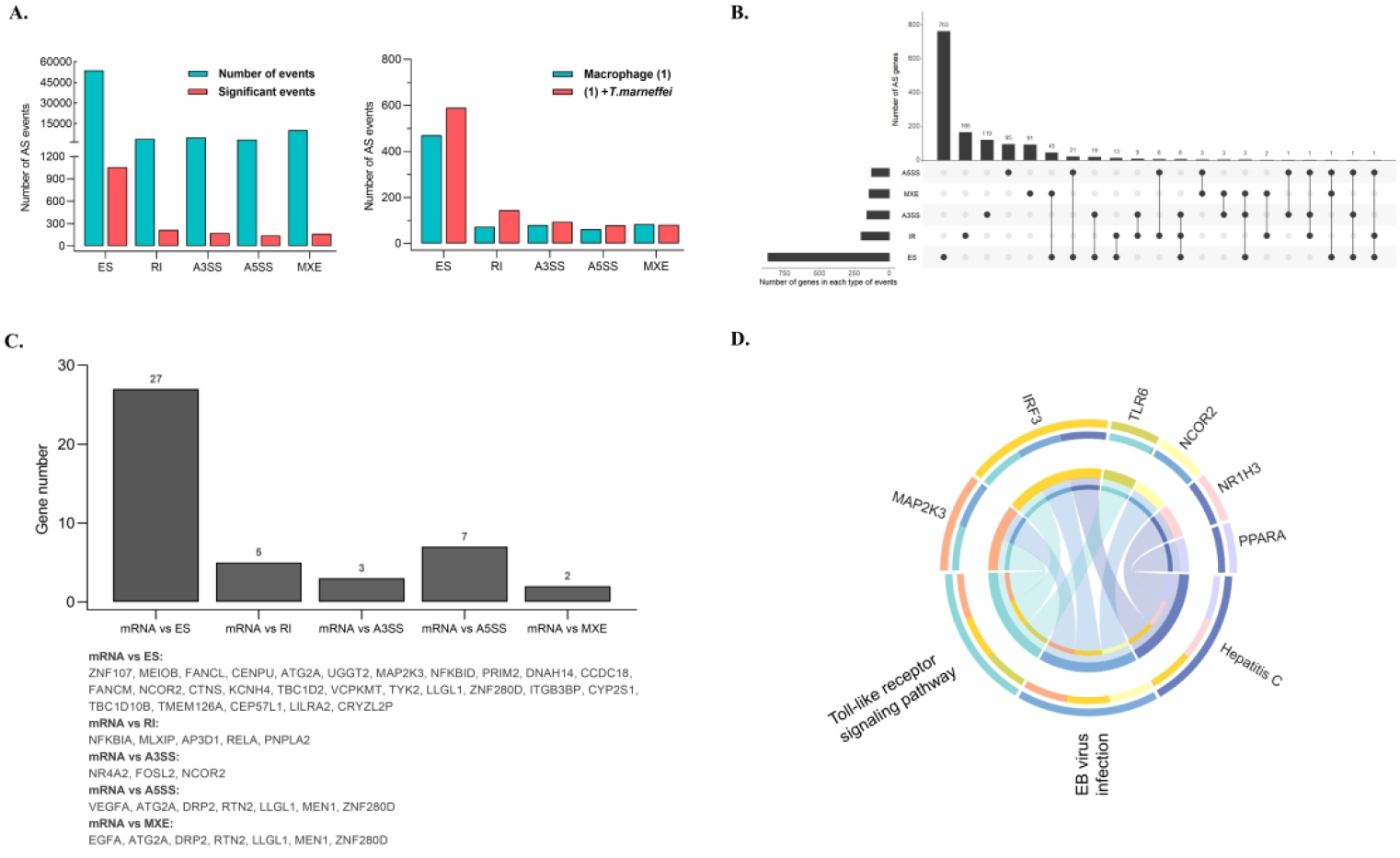
Profiles of alternative splicing in T. marneffei-infected THP-1 macrophages. (A) The number of different types of AS event in T. marneffei-infected macrophages and control cells. (B) UpSet plot (an alternative Venn diagram) of different AS events. (C) Column chart of AS events matched with DEGs which obtained from RNA-seq analysis. (D) Chord diagram of the abundance of top 3 KEGG pathways according to ES-type AS genes after infection of T. marneffei was exhibited.

### *T. marneffei* up-regulated the expression of NCOR2-013 spliceosome in macrophages

Previous studies showed that truncated transcripts play an important role in pathogen immunity (21–23), so we next investigate the influence of AS particularly that of truncated transcript on shaping the cellular response to *T. marneffei* infection. As seen above, *T. marneffei* infection led to an increase in the expression of truncated transcripts of ES-related AS events (NCOR2, NR1H3, PPARA, IRF3, MAP2K3 and TLR6). Considerable differences were observed in terms of the number of reads corresponding to exon-exon junctions between *T. marneffei-infected* group and control group through Sashimi plots and track plots (Fig. 2, A and B), which showed the exon 12 of NCOR2 was spliced, resulting generating NCOR2-013 spliceosome (Fig. 2C, Fig. S4). Subsequently, RNA-seq differential analysis revealed NCOR2-13 spliceosome was significantly up-regulated at 24h *T. marneffei* postinfection (*p*adj < 0.001) (Fig. 2D). Due to the insignificant difference in transcript expression or the inaccuracy of the predicted alternative splicing sites, we only selected NCOR2-013 for in-depth study, the details are shown in Fig. S3, B-J. The qRT-PCR showed that NCOR2-013 spliceosome was increased 2-fold at the mRNA level in *T. marneffei*-infected THP-1 macrophages (Fig. 2E). Additionally, the same trend was exhibited in *T. marneffei*-infected human monocyte-derived macrophage (hMDM), with an approximately 1.5-fold increase (Fig. 2F). Interestingly, the maternal protein of NCOR2, also known as SMRT, was not altered after *T. marneffei* infection, indicating that the biologically functional protein may was not the NCOR2/SMRT, but was more likely to be NCOR2-013 spliceosome (Fig. 2G). These data confirmed that *T. marneffei* leads to the generation of NCOR2-013 spliceosome.

**Figure 2.**
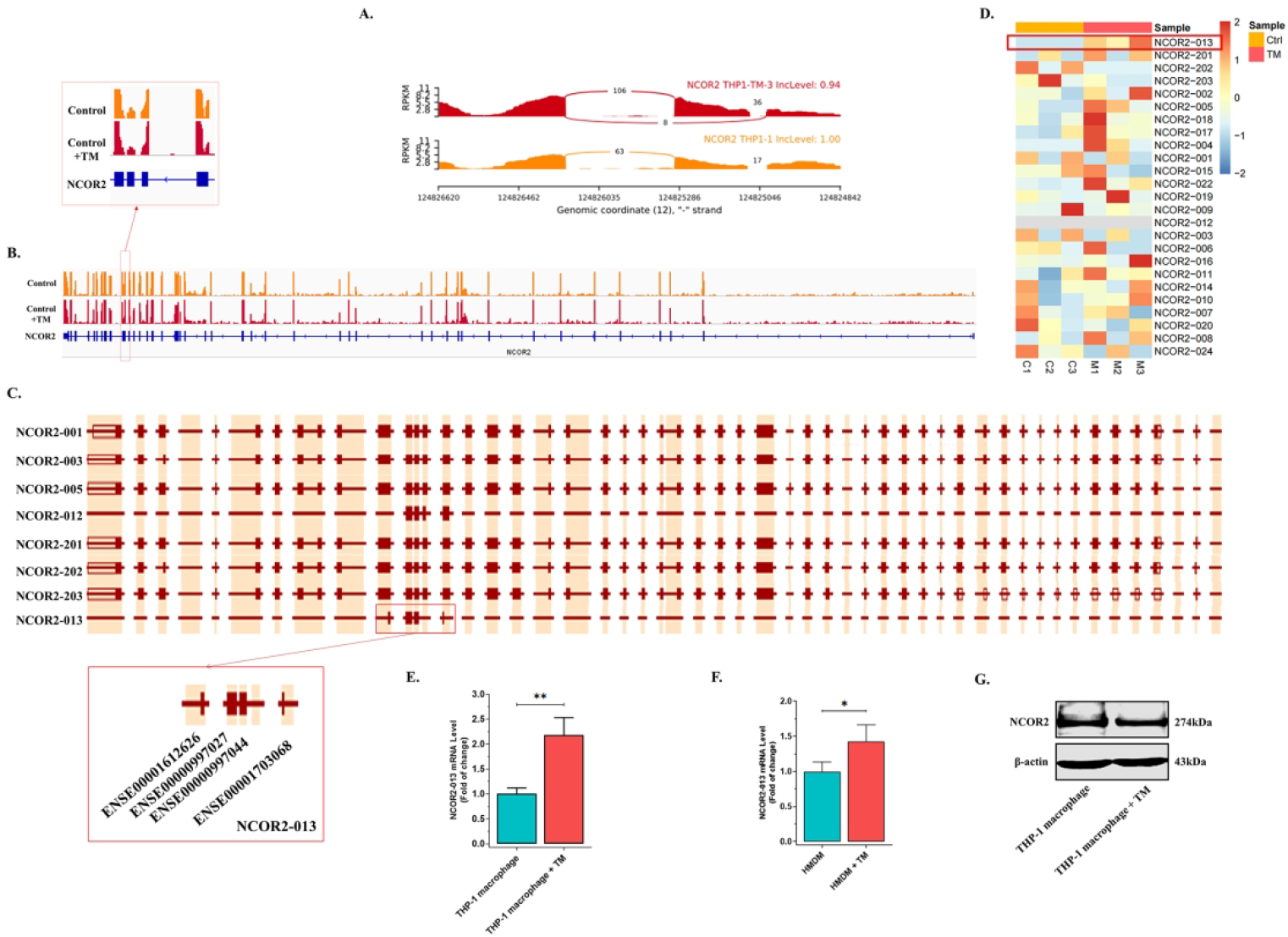
*T. marneffei* up-regulated the expression of the NCOR2-013 spliceosome in macrophages. (A) The Sashimi plot showed the genotype-dependent abundance of splice junctions. The number of observed reads spanning the respective splice junctions is indicated on the Bezier curves, which connect exons. (B) Binding peaks of NCOR2 genes in *T. marneffei*-infected or non-infected macrophages. Gene coding regions were represented asdarker blueannotations at the bottom. The exon and intron were represented by rectangle and straight line, respectively. Exon 1 is on the left. Exon 12 underwent AS of ES-type. The red box represents the site of AS predicted by bioinformatics. (C) The exon-intron organization of NCOR2 transcripts. NCOR2-013 exhibited the missing exon 12, indicating that ES-type AS, induced by *T. marneffei* infection. This plot comes from the public database. (D) The heatmap exhibited the expression levels of different NCOR2 spliceosomes. The red box represents the NCOR2-13 spliceosome was significantly up-regulated after infection with *T. marneffei*. (E-F) qRT-PCR analysis of NCOR2-013 spliceosome expression in THP-1 macrophages and hMDM at 24h *T. marneffei* postinfection. GAPDH was used as a loading control. (G) The WB analysis of NCOR2/SMRT expression at 24h *T. marneffei* postinfection. β-actin was used as a loading control. All data are from three independent experiments, and show mean ± SD. An two-tailed Student’s t-test was used to determine significance, denoted by * (*P* < 0.05), ** (*P* < 0.01), and ns (not significant).

### NCOR2-013 inhibited macrophage inflammatory response and antifungal activity

To validate the NCOR2-013 subcellular localization and function, we successfully constructed the NCOR2-013 overexpression THP-1 macrophage by lentiviruses vectors, validating by qRT-PCR, RNA-seq and WB (Fig. 3, A-C). Subsequently, we used the Hum-mPLoc 2.0 (http://www.csbio.sjtu.edu.cn/bioinf/hum-multi-2/) to predict NCOR2-013 subcellular localization, indicating that the protein mainly perform biological functions in the nucleus (Fig. 3D). Furthermore, we extracted cytoplasm and nucleus protein of *T. marneffei*-infected NCOR2-013 overexpression macrophages and control cells, respectively, and found that *T. marneffei* up-regulated the NCOR2-013 in the nucleus, while hardly detected in the cytoplasm (Fig. 3E). A similar phenomenon was observed in immunofluorescence results (Fig. 3F). Finally, the results of colony forming units (CFU) revealed that more *T. marneffei* were detected in NCOR2-013-overexpression macrophages (Fig. 3G), suggesting that NCOR2-013 inhibited the ability of macrophages to clear *T. marneffei*. The same result was also obtained in hMDM (Fig. 3H). We further explore the effect of NCOR2-013 on the inflammatory response of macrophages, and found that *T. marneffei* up-regulated TNF-ɑ and IL-1β mRNA, while NCOR2-013 overexpression inhibited the elevation of inflammatory factors at mRNA and protein level (NC+*T. marneffei* vs. NCOR2-013 overexpression+*T. marneffei*). Interestingly, *T. marneffei* infection significantly down-regulated TNF-ɑ expression at the protein level, which is the opposite of IL-1β, suggesting *T. marneffei* also regulates TNF-ɑ expression at the post-transcriptional level. All these results indicated that NCOR2-013 overexpression inhibites human macrophage inflammatory response and defense response against *T. marneffei*.

**Figure 3.**
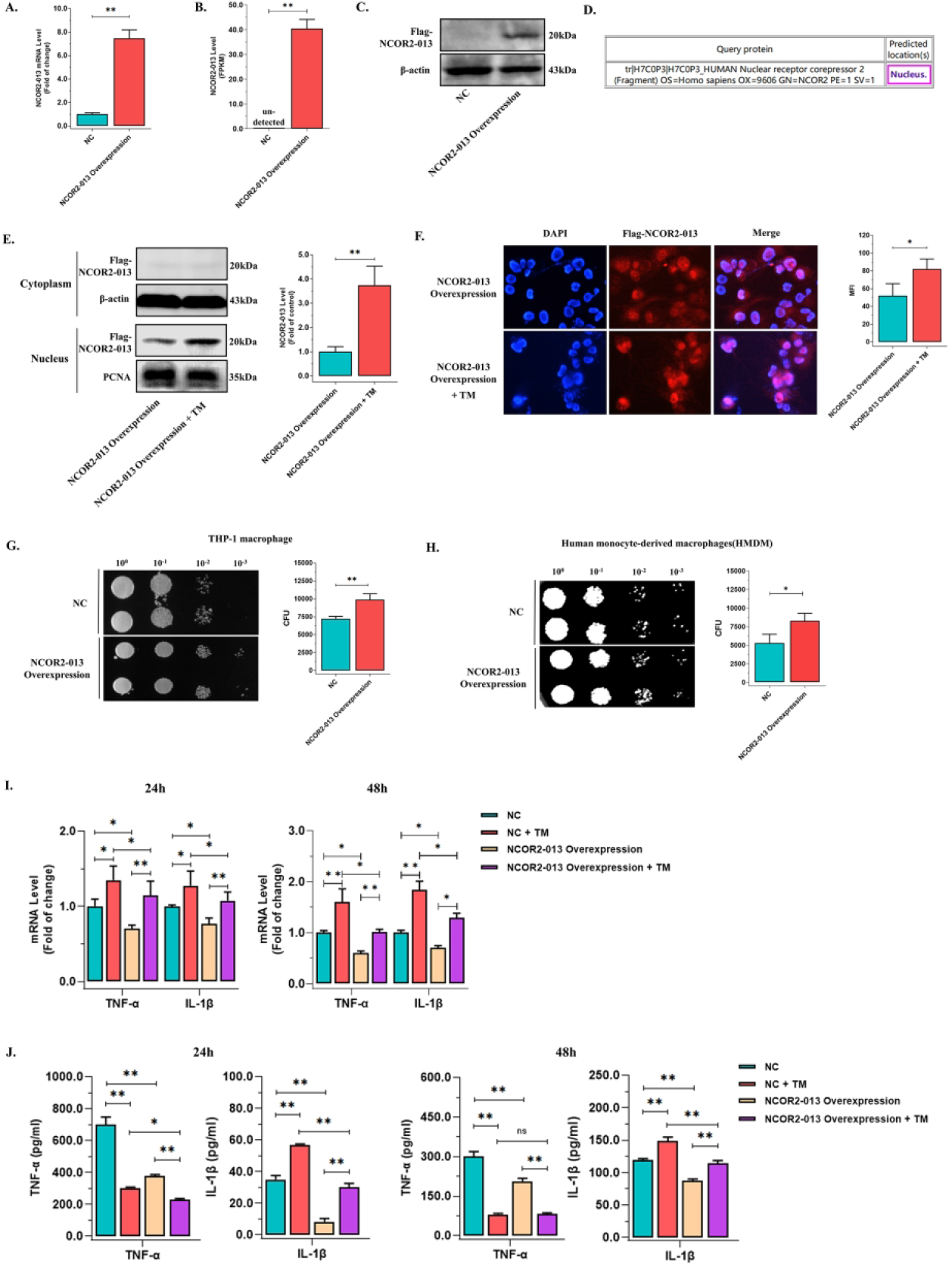
NCOR2-013 inhibited macrophage inflammatory response and antifungal activity. The human NCOR2-013 overexpression THP-1 macrophages or NCOR2-013 overexpression hMDMs were challenged with *T. marneffei* spores (MOI = 10) for 24h or/and 48h. (A-C) NCOR2-013 overexpression confirmed by qRT-PCR (A), RNA-seq analysis (B), and WB (C). (D) Prediction of subcellular localization for NCOR2-013 using the Hum-mPLoc 2.0 (http://www.csbio.sjtu.edu.cn/bioinf/hum-multi-2/). (E) The expression levels of NCOR2-013 in the cytoplasm and nucleus were detected by WB. β-actin and PCNA were used as a loading control for cytoplasm and nucleus, respectively. (F) The subcellular localization of NCOR2-013 was demonstrated by immunofluorescence. Quantification of the data using ImageJ. (G-H) *T. marneffei* colony forming units (CFU) by microdilution spot assay to assess antifungal activity of NCOR2-013 overexpression THP-1 macrophages (G) or hMDMs (H), respectively. (I) The expression levels of TNF-ɑ and IL-1β were detected by qRT-PCR in NCOR2-013 overexpression THP-1 macrophages and control cells infected or non-infected with *T. marneffei*. GAPDH was used as a loading control. (J) The protein levels of TNF-ɑ and IL-1β were measured using CBA human inflammatory cytokines kit in NCOR2-013 overexpression THP-1 macrophages and control cells infected or non-infected with *T. marneffei*. All data are from three independent experiments, and show mean ± SD. An two-tailed Student’s t-test was used to determine significance, denoted by * (*P* < 0.05), ** (*P* < 0.01), and ns (not significant).

### NCOR2-013 suppresses activation of JunB by forming a transcription-regulatory complex with HDAC3 and TBL1XR1/TBLR1

To explore the molecular mechanism by which NCOR2-013 inhibits the inflammatory response, we performed RNA-seq analysis on *T. marneffei*-infected NCOR2-013-overexpression macrophages and control cells, KEGG enrichment analysis showed the DEGs were significantly involved in NF-kappa B signaling pathway and TNF signaling pathway (Fig. S5A), which is consistent with the GSEA results (Fig. S5B), indicating that NCOR2-013 may inhibit the activation of TNF and NF-kappa B signaling pathways.

NCOR2/SMRT is a nuclear receptor that forms a co-repressor with HDAC3 and TBL1XR1/TBLR1 to inhibit inhibit transcription factor activation (24). Since the NCOR2-013 entered the nucleus, we hypothesized that the regulatory mechanism of NCOR2-013 is similar to that of NCOR2/SMRT. Thus, we first identified 8 differentially expressed transcription factors (TWIST1, RB1, STAT1, RELA, HDGF, PPARG, E2F1 and Jun) via TRRUST analysis (25) (Fig. 5, C-D). Then using IP and mass spectrometry to screen for proteins that bound to NCOR2-013 (Fig. 4A). A total of 254 proteins were screened, after intersecting with DEGs, 6 transcription factors were finally obtained (Fig. 4A). Combined with the result of Supplementary Fig. 6D, JunB was finally selected as the most potential transcription factor for subsequent functional studies. To verify hypothesis above, we first verify the protein interaction of NCOR2-013 (Flag-tagged), TBL1XR1\TBLR1, HDAC3, JunB via Co-IP. The result showed that the 4 proteins could bind with each other (Fig. 4, B-E). Next, to investigate whether NCOR2/SMRT was also involved in suppressing JunB, we performed Co-IP to explore the interaction of NCOR2/SMRT with the 4 proteins, interestingly, results showed that NCOR2/SMRT interacts with HDAC3, TBL1XR1/TBLR1, but not NCOR2-013 and JunB (Fig. 4, B-E). Moreover, since both JunD and c-Jun are homologous proteins of JunB (26), we further verified whether NCOR2-013 could bind to them through Co-IP. Interestingly, in addition to JunB, NCOR2-013 was bound by c-Jun, but not by JunD (Fig. 4, F-G). Therefore, we detected the activation levels of JunB and c-Jun by WB, respectively. As a result, the phosphorylation level of JunB showed a significantly lower expression level in NCOR2-013-overexpression macrophages after infection with *T. marneffei* for 24h, while no significant difference was observed for c-Jun (Fig. 4, H-I). Collectively, NCOR2-013 forms the NCOR2-013-HDAC3-TBL1XR1/TBLR1 transcription-regulating complex to inhibit JunB activation.

**Figure 4.**
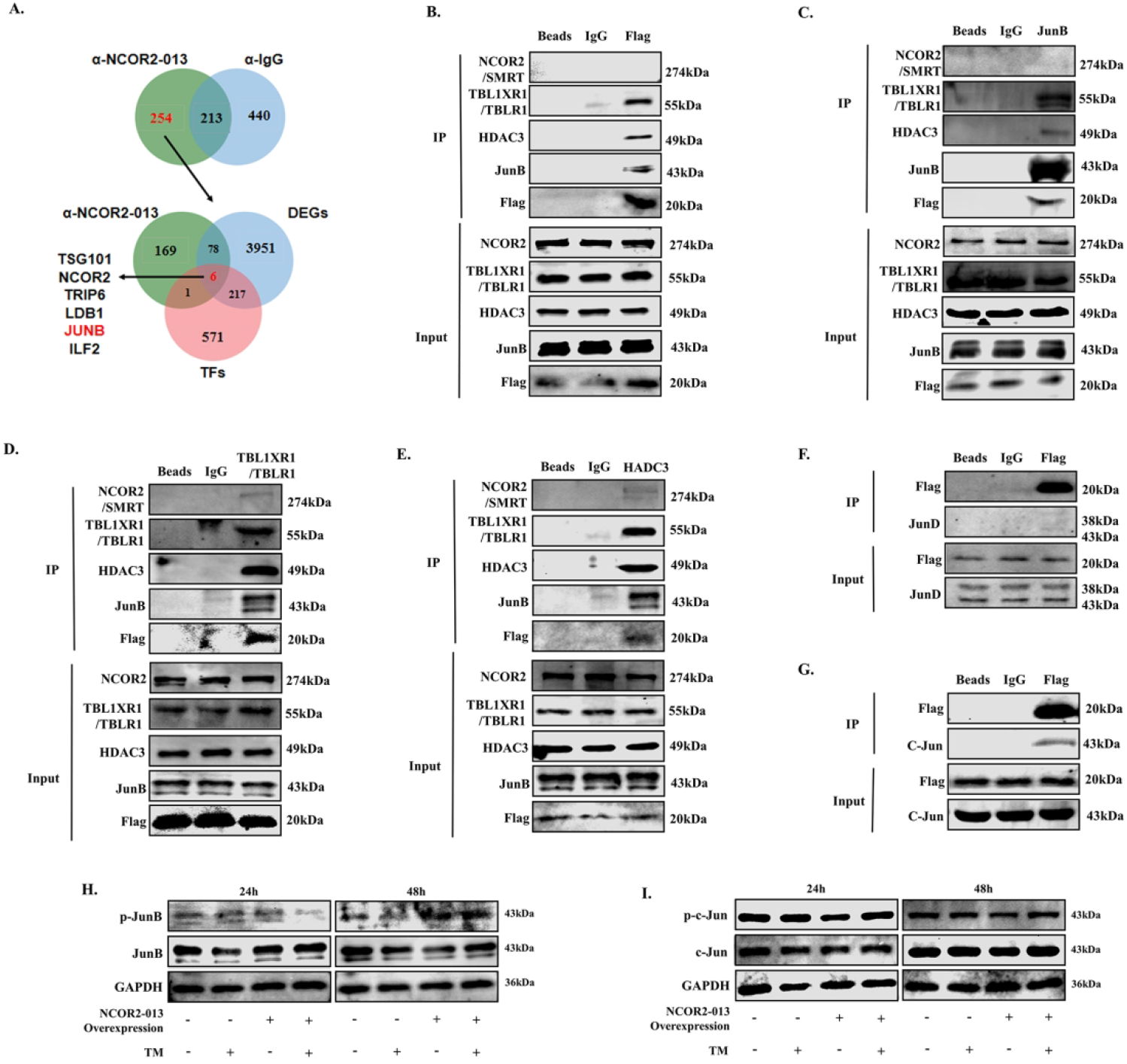
NCOR2-013 suppresses activation of JunB by forming a transcription-regulatory complex with HDAC3 and TBL1XR1/TBLR1. (A) A flowchart of potential NCOR2-013 binding proteins screen. Briefly, NCOR2-013 and isotype control IgG antibodies were used to immunoprecipitate proteins, respectively, and LC-MS was used to detect proteins bound to NCOR2-013. The NCOR2-013 binding proteins were then intersected with DEGs (*T. marneffei*-infected NCRO2-013 overexpression THP-1 macrophages vs. *T. marneffei*-infected control cells) and TFs (list of human transcription factors). (B) IP was performed with Flag-NCOR2-013 antibody and WB was used to detected NCOR2/SMRT, TBL1XR1/TBLR1, HDAC3 and JunB expression. (C-E) IP assays were performed with JunB (C), TBL1XR1/TBLR1 (D), and HDAC3 (E) antibody, respectively, in order to show interactions among TBL1XR1/TBLR1, HDAC3, JunB and NCOR2-013 by WB. (F) Co-IP assays show interactions between Flag-NCOR2-013 and JunD. (G) Co-IP assays show interactions between Flag-NCOR2-013 and c-Jun. (H-I) NCRO2-013 overexpression THP-1 macrophages and control cells were challenged with *T. marneffei* for 24h, 48h, then p-JunB, JunB (H) and p-c-Jun, c-Jun (I) were detected by WB. β-actin was used as a loading control. All data are from three independent experiments.

### NCOR2-013 inhibites pro-inflammatory cytokines transcription by inhibiting acetylated histone H3

To explore whether NCOR2-013 transcription-regulating complex directly regulates transcription of pro-inflammatory cytokines. We performed ChIP-qPCR analysis using NCOR2-013, HDAC3, TBL1XR1/TBLR1 and JunB antibody, respectively, in order to inspect TNF-ɑ and IL-1β expression levels of their promoter regions. The result showed that NCOR2-013, HDAC3, TBL1XR1/TBLR1 and JunB were enriched in the TNF-ɑ and IL-1β promoter regions, indicating that all of the above proteins could act on the promoter region of cytokines (Fig. 5A-B).

**Figure 5.**
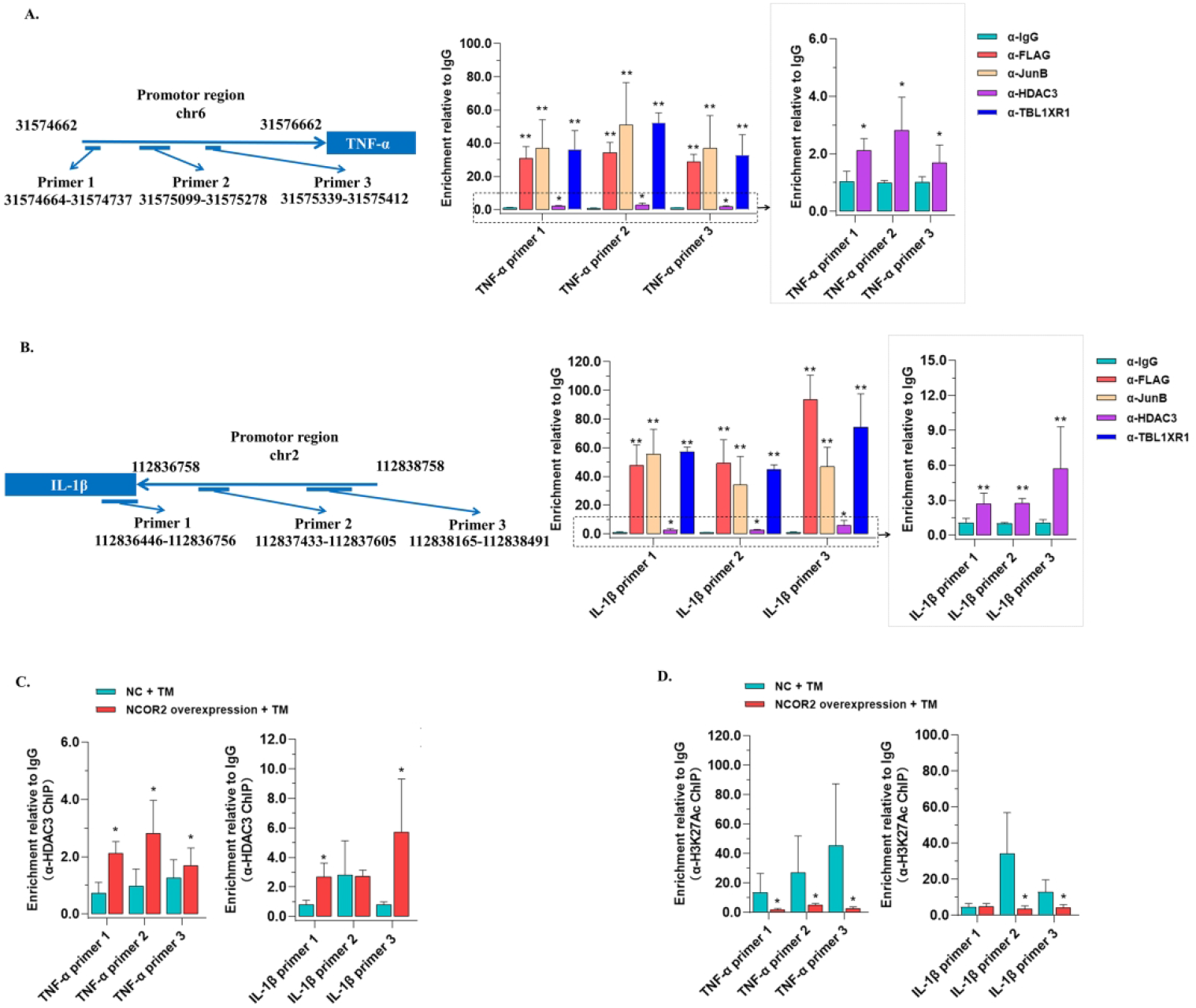
NCOR2-013 inhibites pro-inflammatory cytokines transcription by inhibiting acetylated histone H3. (A-B) The human NCOR2-013 overexpression THP-1 macrophages was challenged with *T. marneffei* spores (MOI = 10) for 24h. ChIP analysis was performed using Flag-NCOR2-013, HDAC3, TBL1XR1/TBLR1, and JunB antibody, respectively, in order to inspect the enrichment levels of these proteins in the TNF-ɑ and IL-1β promoter regions. Three qRT-PCR primers were designed for the promoter region of each cytokine. qRT-PCR assay for promoter regions of TNF-ɑ (A) and IL-1β (B). (C-D) NCRO2-013 overexpression THP-1 macrophages and control cells were challenged with *T. marneffei* for 24h. The enrichment of HDAC3 and histone H3K27Ac in the TNF-ɑ and IL-1β promoter regions of two groups were analysed by ChIP-qPCR. The data was displayed as values of bound/input were relative to IgG. All data are from three independent experiments, and show mean ± SD. An two-tailed Student’s t-test was used to determine significance, denoted by * (*P* < 0.05), ** (*P* < 0.01), and ns (not significant).

In epigenetics, histone acetylation via HDAC3 is a common mechanism that is associated with the transcriptional activation by modulating chromatin condensation (27–29). Thus, to further explore the molecular mechanism of NCOR2-013 suppressing cytokines, NCOR2-013-overexpression macrophages and control cells were infected with *T. marneffei* for 24h, and ChIP-qPCR assays were conducted using HDAC3 and H3K27Ac antibodies. The result showed that NOCR2-013 up-regulated the enrichment level of HDAC3 in promoter regions of TNF-ɑ and IL-1β, and downregulated that of H3K27Ac, indicating that NOCR2-013 inhibited the histone H3 acetylation, which was detrimental for binding of transcriptional factors to the corresponding promoter regions, thereby down-regulating the expression of TNF-ɑ and IL-1β (Fig. 5, C-D).

### LPS, but not JunB overexpression or IFN-γ, blocks inhibition of pro-inflammatory response by NCOR2-013 in macrophage

To explore how to reverse the inhibition of pro-inflammatory responses by NCOR2-013, we overexpressed JunB in NCOR2-013-overexpression THP-1 macrophages, however, it hardly reverse the inhibitory effect of NCOR2-013, evidenced by the fact that the JunB-overexpression group did not have significant changes in cytokine expression and antifungal ability (Fig. S6, A-C). Subsequently, we conducted exploration experiments using IFN-γ and LPS, respectively. Interestingly, the LPS, but not IFN-γ (Fig. S6, D-E), was able to significantly upregulate the expression of TNF-ɑ and IL-1β, while significantly increasing the killing ability of macrophages against *T. marneffei* (Fig. S6, F-G). Collectively, these data suggest that LPS, but not the JunB overexpression or IFN-γ, reverses the NCOR2-013-mediated inhibition of pro-inflammatory responses in macrophages.

### Identification of TUT1, which generates NCOR2-013 spliceosome by regulating AS, as an upregulated gene closely associated with *T. marneffei* immune evasion

To identify the upstream RBP that regulate NCOR2 for AS, we predicted splicing regulators using rMATs and then intersected them with DEGs obtained by RNA-seq analysis, the result showed that TUT1 might be the RBP with statistical significance (Fig. 6A), with the motif was [AC][AG]ATACT (Fig. 6B). qRT-PCR and WB results showed that *T. marneffei* up-regulated the expression of TUT1 in THP-1 macrophages (Fig. 6C). Afterwards, we performed the visualization of origin genome profiles via IGV, showing that TUT1 motif could be combined with exon 12 of NCOR2 (Fig. 6D), which was consistent with the above-mentioned (Fig. 2). To determine whether the effect on NCOR2 AS was mediated by direct binding of TUT1, we performed RIP and found that TUT1 binding to NCOR2 pre-mRNA (Fig. 6E). Moreover, the expression of NCOR2-013 isofrom was significantly increased in the *T. marneffei-*TUT1-overexpression THP-1 macrophages, while the expression of NCOR2-013 isofrom was significantly decreased in the *T. marneffei*-TUT1-knockdown THP-1 macrophages (Fig. 6, F-I). These finding indicats that TUT1 performed an important regulatory function in the occurrence of AS in NCOR2 after infection of *T. marneffei*.

**Figure 6.**
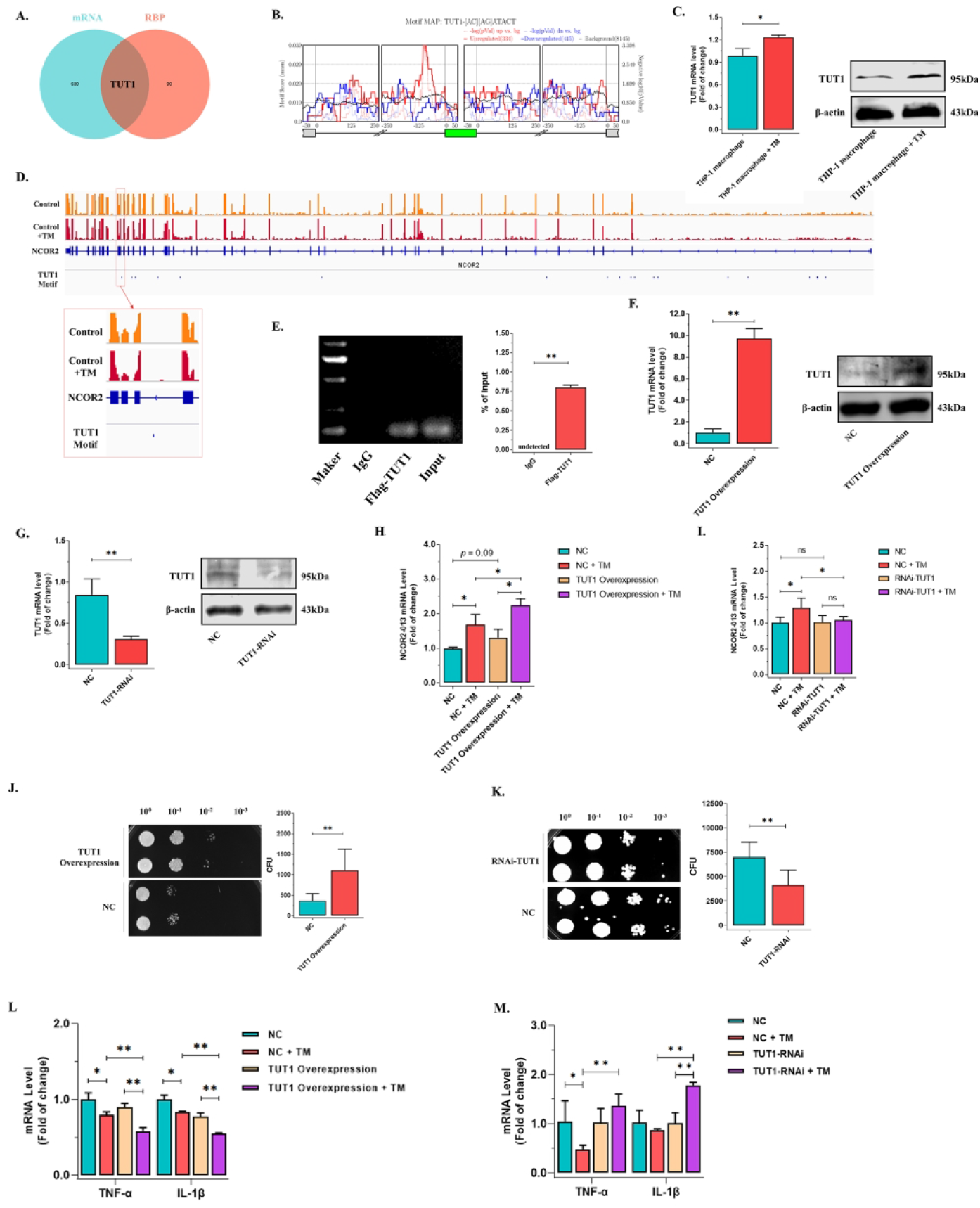
TUT1-mediated generation of the NCOR2-013 spliceosome to suppress inflammatory in *T. marneffei*-infected THP-1 macrophages. (A) flowchart to identify potential RBP that regulating AS events. Briefly, rMATs was used to predict potential RBPs and then were intersected with DEGs. (B) The motif map of TUT1 that prediction via rMATs. (C) qRT-PCR and WB to detect the expression level of TUT1 in *T. marneffei*-infected or non-infected THP-1 macrophages. (D) Visualization of origin genome profiles was performed with IGV. TUT1 motif could be combined with exon 12 of NCOR2. (E) RIP analysis to verify the interaction of TUT1 and NCOR2 pre-mRNA, a Flag-tagged TUT1 was included as an indicator when constructing TUT1-overexpressing macrophages. IP was performed with Flag antibody and RNA was subjected to agarose gel electrophoresis with subsequent RNA gel blot analysis. (F-G) TUT1-overexpression (F) and TUT1-knockdown (G) in THP-1 macrophage confirmed by qPCR and WB. (H-I) The expression of NCOR2-013 was detected by qRT-PCR in *T. marneffei*-infected or non-infected TUT1-overexpression THP-1 macrophages (H) and TUT1-knockdown THP-1 macrophages (I). (J-K) *T. marneffei* CFU by microdilution spot assay to assess antifungal activity of TUT1-overexpression THP-1 macrophages (J) and TUT1-knockdown THP-1 macrophages (K). (L-M) The expression of TNF-ɑ and IL-1β were detected by qRT-PCR in TUT1-overexpression/knockdown THP-1 macrophages and control cells infected or non-infected with *T. marneffei*. GAPDH was used as a loading control. All data are from three independent experiments, and show mean ± SD. An two-tailed Student’s t-test was used to determine significance, denoted by * (*P* < 0.05), ** (*P* < 0.01), and ns (not significant).

We then investigated the effect of TUT1 on *T. marneffei* clearance in THP-1 macrophages. As a result, more *T. marneffei* were detected in TUT1-overexpression THP-1 macrophages (Fig. 6J), and the inflammatory factors (TNF-α, IL-1β) were down-regulated (Fig. 6L), but opposite trend was observed in TUT1-knockdown macrophages (Fig. 6, K and M), suggesting that *T. marneffei*-mediated TUT1 impaired macrophage clearance of fungi. Collectively, these data suggest that *T. marneffei* suppresses macrophages inflammatory by producing the truncated protein NCOR2-013 via TUT1-regulated AS (Fig. 7).

**Figure 7.**
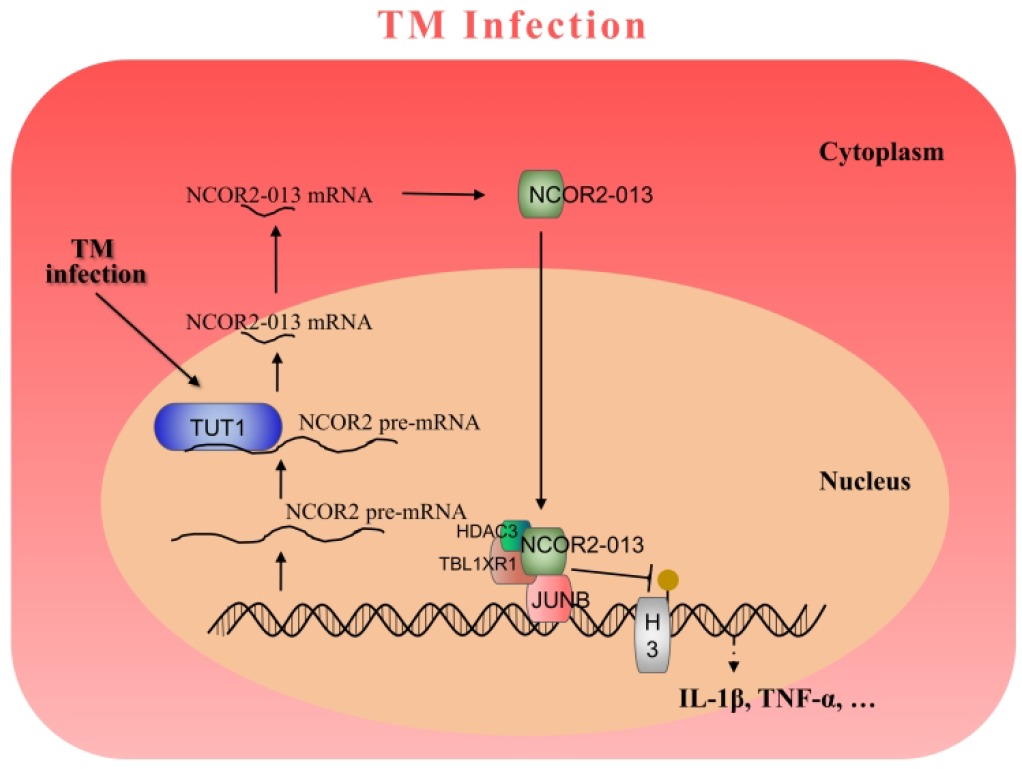
*T. marneffei* suppresses macrophages inflammatory by producing the truncated protein NCOR2-013 via TUT1-regulated AS. 486×329mm (291 × 291 DPI)

## DISCUSSION

The diverse products resulting from alternative splicing is attributed to play a crucial role in the biological functions, especially for the the regulation of cellular immune response, is gradually getting evident (30,31). In this study, we reported a sharp change in the macrophage gene AS events profile upon *T*.*marneffei* infection by high-throughput RNA sequencing. Furthermore, we found that NCOR2-013, a truncated protein of NCOR2/SMRT, was significantly upregulated during *T. marneffei* infection. Mechanistically, NCOR2-013 can inhibit JunB-mediated transcriptional activation of pro-inflammatory factors, TNF-α and IL-1β. Also, we identified a AS regulator RBP, TUT1 that involved in facilitating *T*.*marneffei* immune evasion via regulation of NCOR2-013 production. Taken together, the results indicate that *T*.*marneffei* escapes macrophages killing through the TUT1-mediated the alternative splicing of NCOR2-013.

Innate immunity is the first line of defense against pathogens invasion, However, several evasion strategies against the host’s innate immunity have gradually evolved during pathogens evolution. Among them, the most common strategy is to inhibit the activation of innate immune pathways, such as TLR pathway, cGAS-STING pathway, IFN pathway, by interacting with host protein through pathogen virulence protein (10,32–34). In recent years, more and more studies have found that AS plays an important role in regulating innate immunity due to the rapid development of high-throughput sequencing technologies (30,31). Thousands of AS events have been detected in human dendritic cells and macrophages induced by bacterial challenge (22,35,36). So far, little is known about the landscape of AS events associated with *T*.*marneffei*-infected macrophages. Thus, we have conducted comprehensive repertoire of AS events in human THP-1 macrophage challenged with *T*.*marneffei* and obtained several significant finding. Firstly, we identified differentially 1,757 AS events in *T*.*marneffei*-infected macrophages compared to control, which is similar to other bacterial, like Mtb (22), suggesting gene splicing augments occurred upon pathogen-host interaction. Subsequently, the genes that harbor differential AS events were highly enriched for the pathogen immune-related signaling pathways, indicating splicing of these genes may serve as another mechanism for macrophages resist *T*.*marneffei* or *T*.*marneffei* immune escape. Finally, 6 master regulators of *T*.*marneffei* immune-related signaling pathways, NCOR2, NR1H3, PPARA, IRF3, MAP2K3 and TLR6, were identified that have the most significant differentially AS events, suggesting AS are involved in the core link of regulating *T*.*marneffei* immune response.

Typically, protein-coded genes can produce alternatively spliced variants of multiple transcripts, including some non-functional and/or truncated variants of the protein, that mediate different biological functions. The reasons for the significant biological differences of these alternately spliced variants may include that the truncated regions have certain biological functions, such as signal peptides, DNA binding sites, post-translational modification sites, etc. In immune cells, the expression levels of different alternative spliced variants may have a significant impact on activation of signaling pathways, immune status, and antibacterial capacity. For instance, MyD88L and MyD88S is a long isoform and short isoform of MyD88 protein, which play a role in activating and inhibiting the innate immunity, respectively (37,38). Dengue virus deregulating innate immune response via inducing SAT1 exon 4 skipping to produce an isoform of SAT1 (21). Also, the up-regulation of RAB8B truncated variant helps Mtb evade killing from macrophage phagosome (22). However, alternatively spliced variants have been poorly studied in the field of fungal-host interaction.

Here we identified an alternative spliced variant, NCOR2-013, in the *T*.*marneffei*-infected macrophages. Notably, normal THP-1 macrophages hardly express NCOR2-013 (Figure 3), suggesting this isoform may not be the protein that maintains the normal metabolism of cells. Thus, *T*.*marneffei* may use this isoform to implement some biological process that is beneficial to its own survival in macrophage. NCOR2 (also known as SMRT) is a large nuclear receptor co-repressor present in the nucleus with a molecular weight of up to 274 kDa (24). Interestingly, as one of the 25 isoforms of NCOR2/SMRT, the molecular weight of NCOR2-013 is 17kDa, only one-sixteenth of NCOR2/SMRT (39,40). Structurally, compared with NCOR2/SRMT, NCOR2-013 only retains 4 exons. Nevertheless, NCOR2-013 still retains a biological function similar to that of NCOR2/SMRT, acting as part of a multisubunit complex which includes TBL1XR1/TBLR1 and HDAC3 to inhibit chromatin unfolding that specifically blocks JunB-mediated basal transcriptional activity of pro-inflammatory genes (24,41,42). The function of the *T*.*marneffei*-induced short isoform is similar to long isoform, which is not the same as the other pathogens mentioned above (22,35,36). Our data demonstrate that overexpression of NCOR2-013 significantly inhibited macrophage inflammatory response and its antifungal activity, suggesting that it has a highly effective anti-inflammatory function, considering its small molecular weight. Our Co-IP results showed that NCOR-013 did not bind to NCOR2/SMRT, suggesting that NCOR-013 exerts anti-inflammatory effects independently of NCOR2/SMRT. Furthermore, we found that the binding affinity of JunB to NCOR2-013 is significantly higher than that of NCOR2/SMRT, indicating *T*.*marneffei* may use NCOR2-013 instead of NCOR2/SMRT to effectively suppress inflammatory responses. More interestingly, even overexpression of JunB did not reverse the anti-inflammatory response of NCOR2-013, which may implying the strong affinity of NCOR2-013 for JunB. Moreover, the LPS, but not IFN-γ, blocks NCOR2-013-mediated inhibition of pro-inflammatory responses in macrophages, suggesting that NCOR2-013 may only mediate inhibition of specific inflammatory pathways. Previous studies have shown that activation of the IFN-γ pathway relieved the inhibitory effect of NCRO2/SMRT, while activation of the TLR4 pathway can relieve the inhibitory effect of NCOR (24). Thus, from our results, the biological role of NCOR2-013 may be similar to that of NCOR, but not to that of NCRO2/SMRT. Further studies need to be conducted to reveal the specific molecular mechanism that causes this specific activation.

From the perspective of epigenetic mechanism, the NCOR2-013 regulatory complex may depend on HDAC3 to inhibit histone acetylation modification, thereby inhibiting the transcriptional activation response of pro-inflammatory factors (24). HDAC3 is a chromatin-modifying enzymes that silence transcription via the inhibition of histones acetylation. Traditionally, HDAC3 need to interaction with NCoR2/SMRT to engage its catalytic activity (43–45). Interestingly, we found that HDAC3 can interaction with NCOR2-013 instead of NCOR2/SMRT, thereby exerting an inhibitory effect, which similar to the results of a recent study, that is HDAC3 is recruited to ATF2-bound sites without NCOR2/SMRT (46). In this study, we found that compared to the control, *T*.*marneffei*-infected NCOR2-013-overexpression macrophages had more HDAC3 protein enrichment in pro-inflammatory promoter regions, and conversely, histone acetylation levels in these regions were significantly reduced. Similar to other results, these findings demonstrate that HDAC3 mediates macrophage reactivity in inflammatory macrophages (47). Taken together, we speculate that *T*.*marneffei* may use NCOR2-013 to efficiently achieve immune evasion.

AS processes are mainly controlled by splicing regulators, among which are RBPs. Abnormal splicing regulators expression often elicit changes in alternative splicing to promote or inhibit the antipathogenic activity of immune cells (30,31). SRSF1 and PTBP1 promotes the exclusion or inclusion of exon 13 in CD46, which involved in pathogen infection (48).Meanwhile, RBM10 is responsible for SAT1 exon 4 skipping for limiting dengue viral replication (21).In this study, TUT1 was identified as a anti-inflammatory gene that was important for NCOR2-013 production and *T*.*marneffei* immune evasion. TUT1 is a nucleotidyl transferase that functions as both a terminal uridylyltransferase and a nuclear poly(A) polymerase, which specifically adds and removes nucleotides from the 3’ end of small nuclear RNAs and select mRNAs, affecting gene expression (49,50). TUT1 associates with RBM10 to control the expression and 3’ end processing of cardiac mRNAs, suggesting it may act as an alternative splicing regulator (51). However, little is known about the role of TUT1-mediated alternative splicing in immune responses. In this study, we identified a novel transcript NCOR2-013 that was produced in *T*.*marneffei*-infected macrophages with TUT1 upregulation. TUT1 overexpression reduced the expression of NCOR2-013 and pro-inflammatory factors, so as the antibacterial capacity of macrophages, in part by regulating alternative splicing of NCOR2, and conversely, TUT1 knockdown showed the opposite phenomenon, suggesting that TUT1 acts as a splicing factor that promotes *T*.*marneffei* immune escape.

In summary, we have systematically analyzed alternative splicing events in *T*.*marneffei*-infected macrophages and identified TUT1 as the splicing regulator that was important for the NCOR2-013 production and *T*.*marneffei* immune evasion. TUT1 inhibits inflammatory response via inducing AS process of NCOR2 to produce the novel transcript NCOR2-013. This truncated protein NCOR2-013 acts as an important immune escape-related factor for macrophages through recruiting TBL1XR1/TBLR1 and HDAC3 to form a regulatory complex to inhibits histone acetylation modification, which finally specifically blocks JunB-mediated basal transcriptional activity of pro-inflammatory genes (Figure 7). TUT1 and NCOR2-013 represent the potential targets for the development of mechanism-based talaromycosis prevention strategies. However, some issues are still unclear and need further research in the future. First, results found that NCOR2/SMRT still forms a complex with HDAC3 and TBL1XR1/TBLR1, so what role does it play in *T*.*marneffei* infection? Second, the exact mechanism by which NCOR2-013 forms the regulatory complex is unclear. Last, how *T*.*marneffei* infection affects the expression of TUT1 still needs to be systematically studied.

## DATA AVAILABILITY

We deposited the raw fastq files in the Sequence Read Archives (SRA) of the National Center for Biotechnology Information (NCBI) under accession number GSE200512, GSE154779 of Bioproject PRJNA824858, PRJNA647412, respectively.

## SUPPLEMENTARY DATA

Supplementary Data are available at NAR online.

## ACKNOWLEDGEMENTS

Hao Liang, Li Ye, Junjun Jiang, Wudi Wei, and Gang Wang designed experiments and provided conceptual input. Chuanyi Ning, Jingzhen Lai, Zongxiang Yuan and Rongfeng Chen provided reagents. Wudi Wei, Hong Zhang, Xiuli Bao, Jinhao He, Lixiang Chen, and Yuxuan Liu performed experiments. Wudi Wei, Gang Wang, Hong Zhang, Sanqi An and Qiang Luo analyzed data. Wudi Wei and Gang Wang wrote the manuscript.

## FUNDING

The study was supported by National Natural Science Foundation of China (NSFC; 81971934), Guangxi Bagui Scholar (to Junjun Jiang), Thousands of Young and Middle-aged Key Teachers Training Program in Guangxi Colleges and Universities (to Junjun Jiang), Guangxi Medical University Training Program for Distinguished Young Scholars (to Junjun Jiang), China Postdoctoral Science Foundation (2020M683212, to Wudi Wei), Guangxi Postdoctoral Special Foundation (to Wudi Wei) and Guangxi Youth Science Fund Project (2021GXNSFBA196004, to Wudi Wei).

## Conflict of interest statement

None declared.

## Supplemental material

**Fig. S1.**
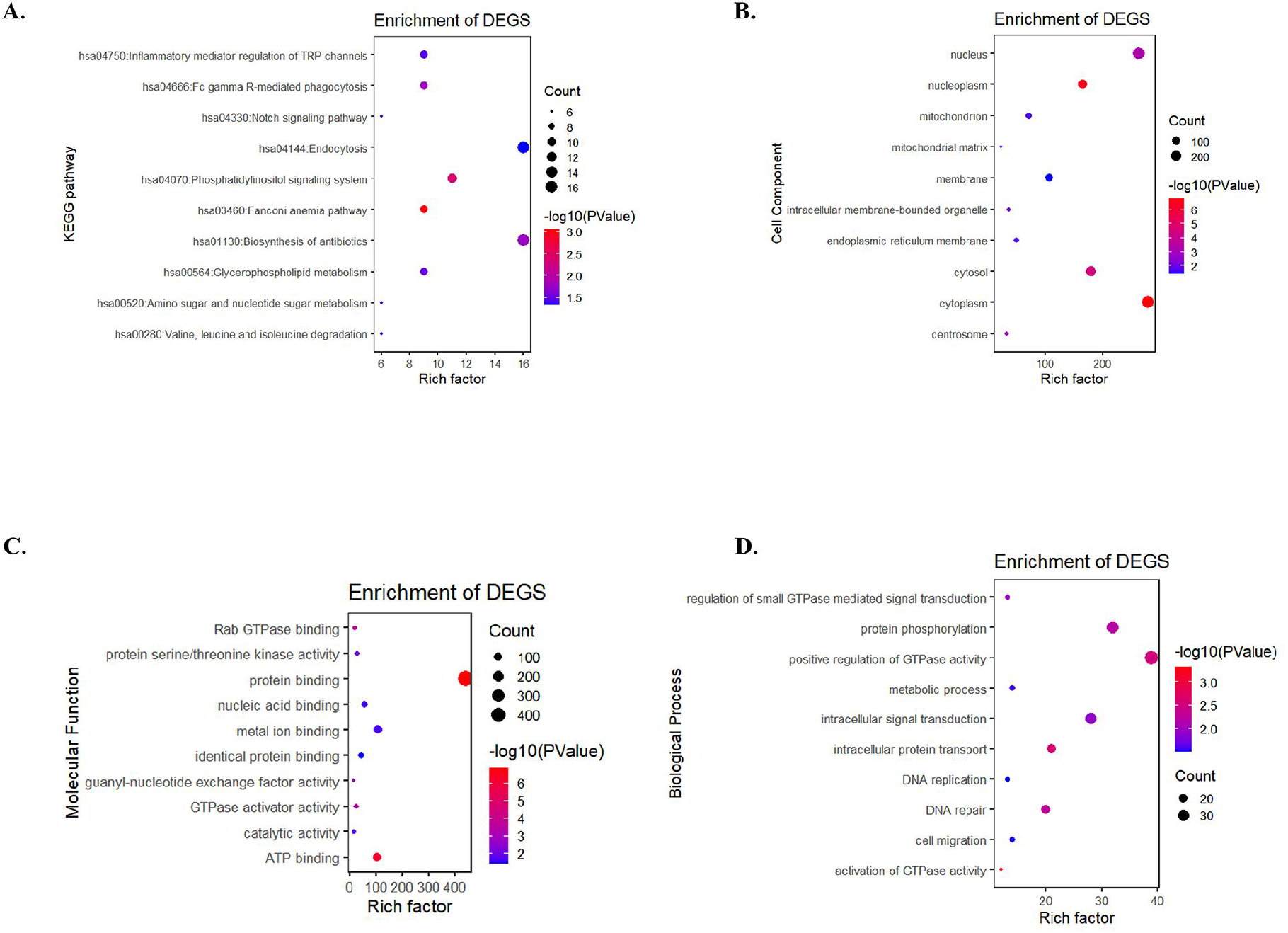
GO and KEGG pathway functional enrichment analysis of ES-type AS events.

**Fig. S2.**
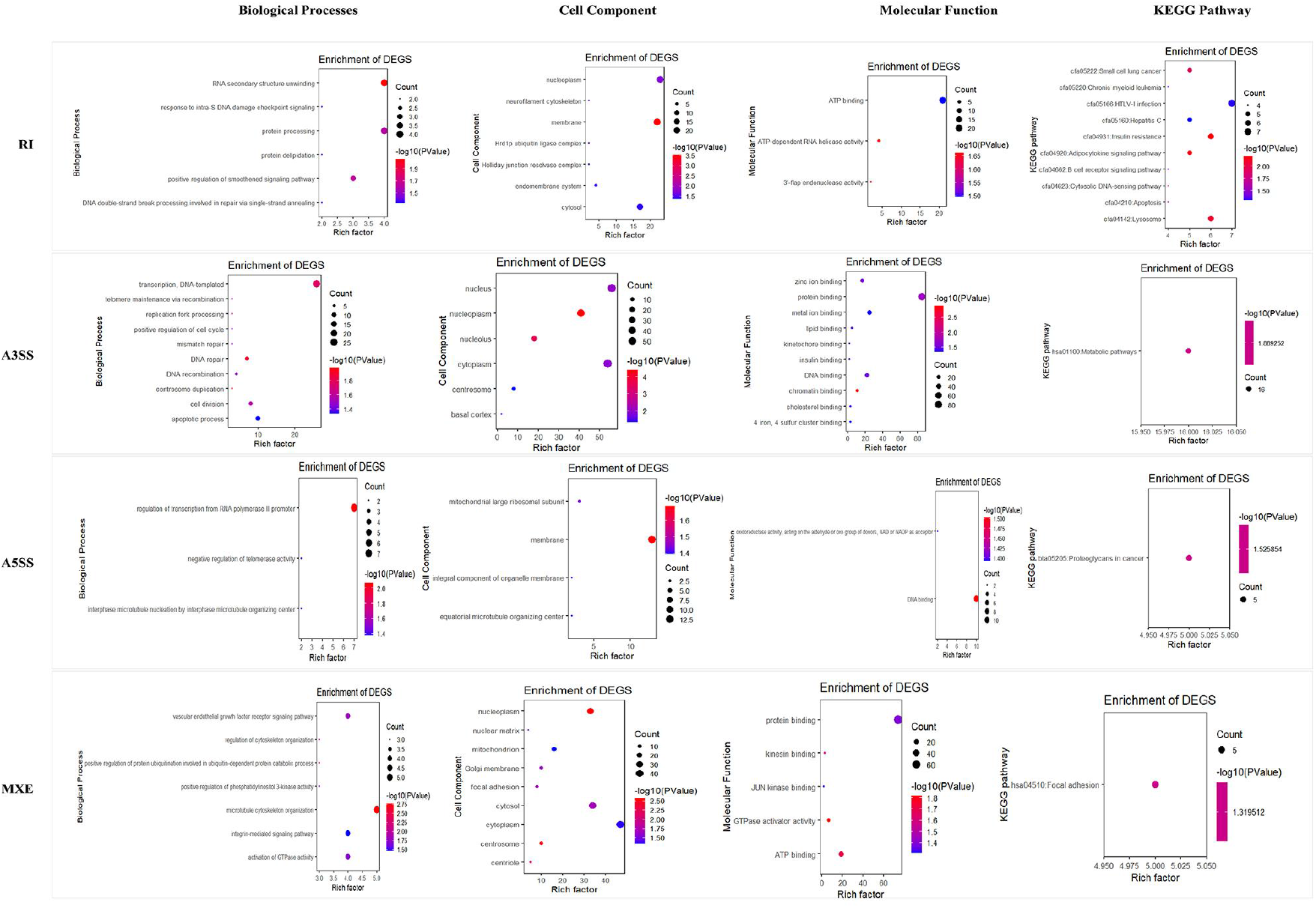
GO and KEGG pathway functional enrichment analysis of other AS event types, RI, A3SS, A5SS and MXE.

**Fig. S3.**
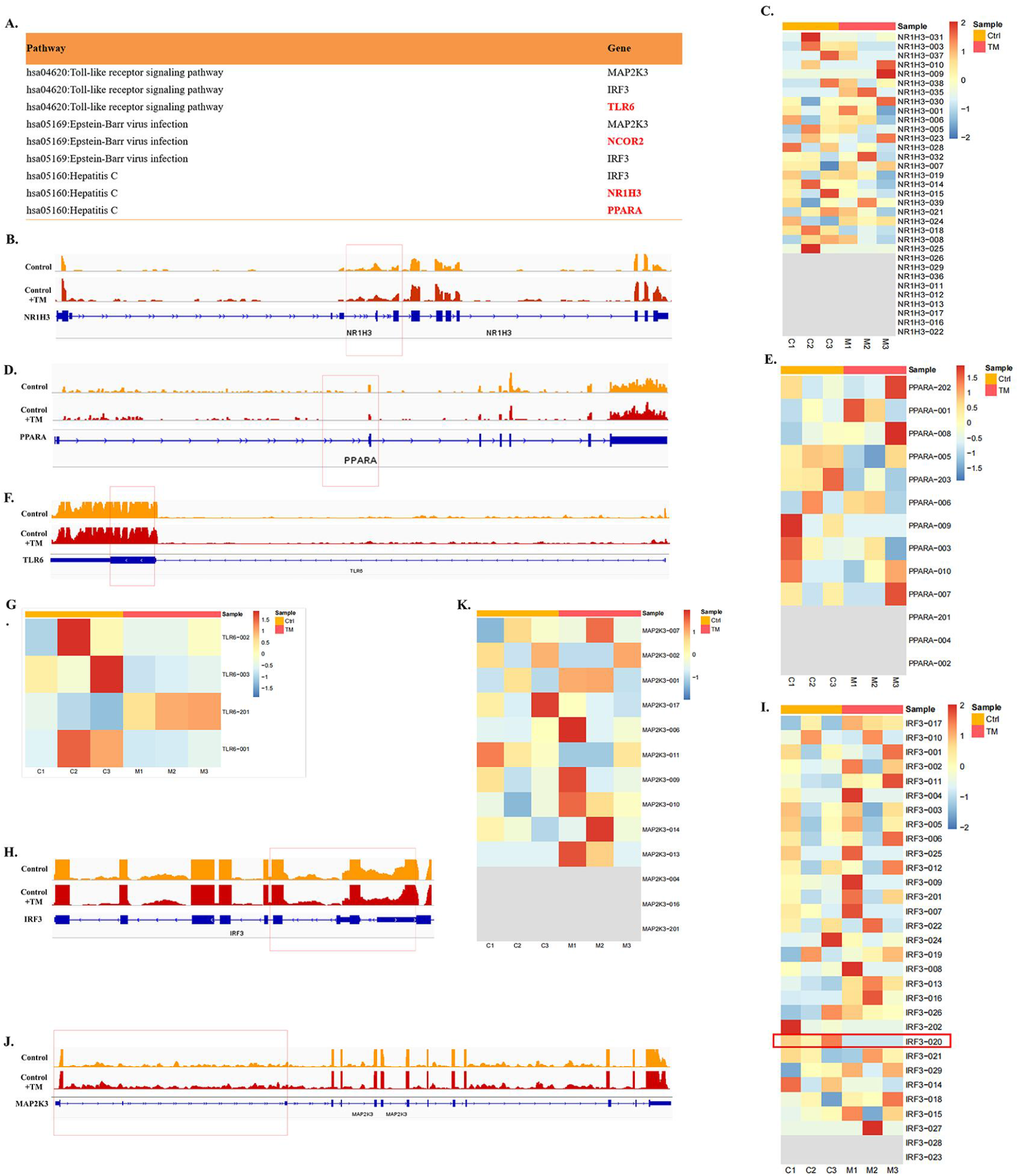
The heatmaps and track plots of NR1H3, PPARA, IRF3, MAP2K3 and TLR6 transcripts. (A) Summary of the abundance of top 3 KEGG pathways according to ES-type AS genes, with FDR < 0.05 and |Lnclevel| > 0.5, at 24h T. marneffei postinfection was exhibited. (B, D, E, H, J) Binding peaks of NR1H3, PPARA, TLR6, IRF3, MAP2K3 genes in *T. marneffei*-infected or non-infected macrophages. Gene coding regions were represented asdarker blueannotations at the bottom. The exon and intron were represented by rectangle and straight line, respectively. The red box represents the site of AS predicted by bioinformatics. (C, E, G, K, I) The heatmap exhibited the expression levels of different NR1H3, PPARA, TLR6, IRF3, MAP2K3 spliceosomes. It is worth mentioning that there were also differential transcripts in NR1H3, PPARA and IRF3, but showed a decreasing trend, which was inconsistent with the predicted results of bioinformatics. Although one of TLR6 transcripts showed an increasing trend in *T. marneffei*-infected macrophages, the P value was greater than the screening threshold (padj < 0.001). Comprehensive considering, the above four transcripts were filtered out.

**Fig. S4.**
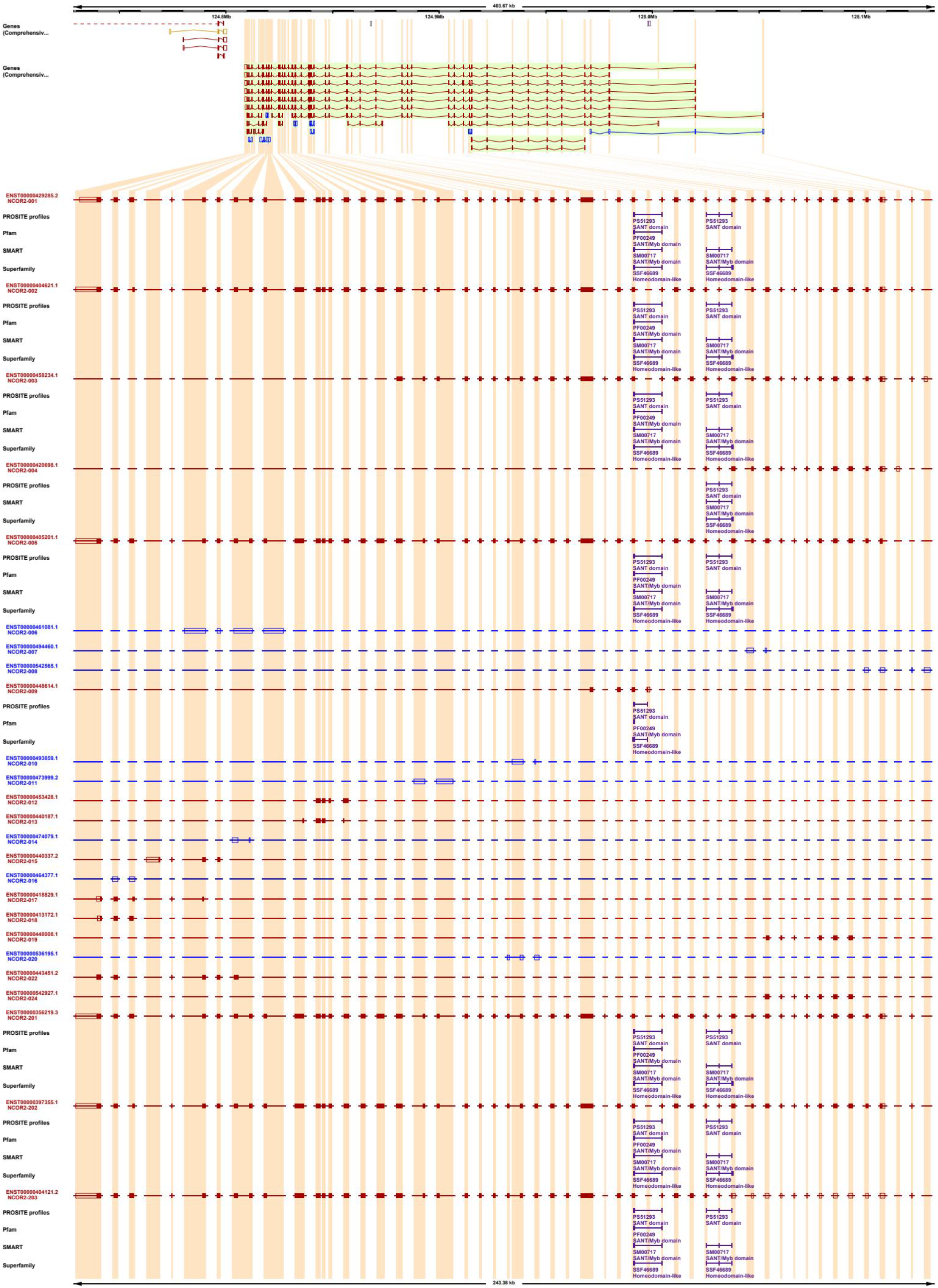
The complete exon-intron organization of NCOR2 transcripts.

**Fig. S5.**
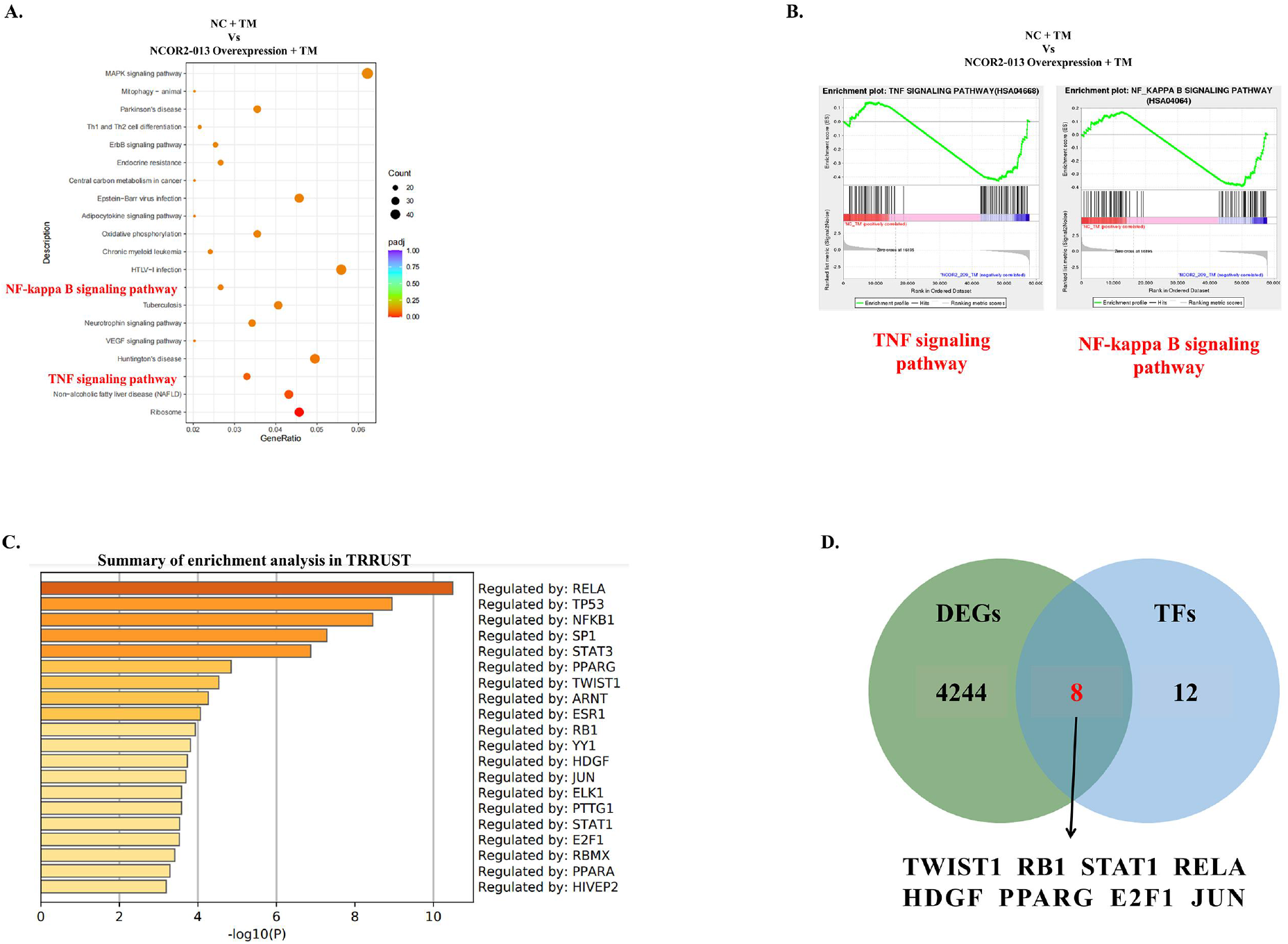
Profiles of transcriptome level of NCRO2-013 overexpression THP-1 macrophages infected with *T. marneffei*. (A-B) KEGG pathway functional (A) and GSEA (B) enrichment analysis of DEGs in T. marneffei-infected NCRO2-013 overexpression THP-1 macrophage conpared with control. (D)TRRUST analysis was performed to predict the set of transcription factors regulating differential genes. (E) The predicted 20 transcription factors were intersected with DEGs, and finally 8 transcription factors that may play a role in the regulation of T. marneffei infection by NCOR2-013 were obtained, namely TWIST1, RB1, STAT1, RELA, HDGF, PPARG, E2F1 and JUN.

**Fig. S6.**
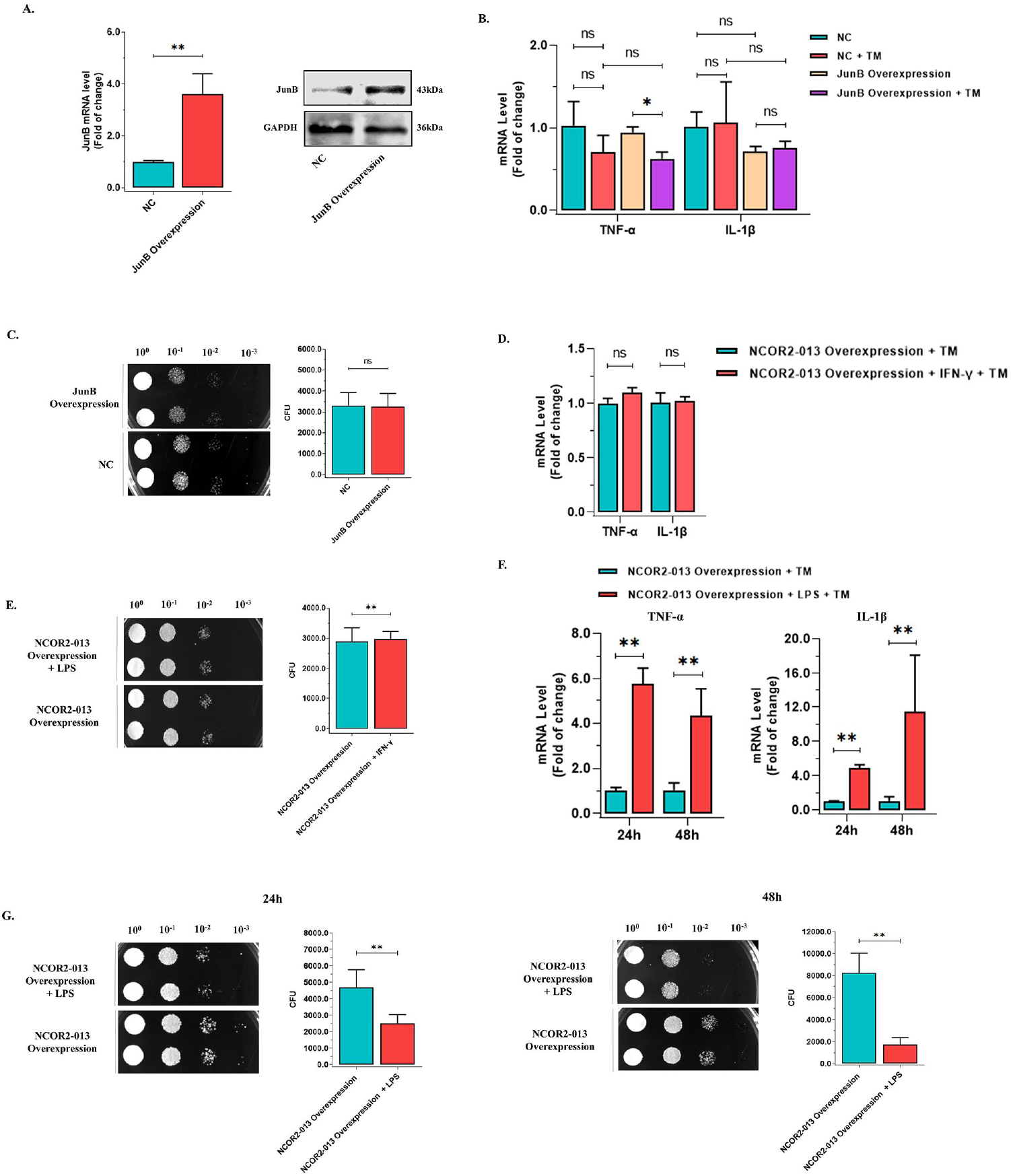
LPS, but not JunB overexpression or IFN-γ, blocks NCOR2-013-mediated inhibition of pro-inflammatory responses in macrophages. (A) JunB overexpression in NCOR2-013 overexpression THP-1 macrophages confirmed by qRT-PCR and WB. (B) The expression of TNF-ɑ and IL-1β was detected by qRT-PCR in *T. marneffei*-infected or non-infected JunB-overexpression THP-1 macrophages and conrol cells. (C) *T. marneffei* CFU by microdilution spot assay to assess antifungal activity of JunB-overexpression THP-1 macrophages. (D-E) The NCOR2-013 overexpression THP-1 macrophages and control cells were stimulated with IFN-γ (100ng/ml) for 24h, and were infected or non-infected with *T. marneffei* spores (MOI = 10) for 24h. (D) qRT-PCR assays for TNF-ɑ and IL-1β expression in IFN-γ-stimulated NCOR2-013 overexpression THP-1 macrophages. (E) *T. marneffei* CFU by microdilution spot assay to evaluate antifungal activity of IFN-γ-stimulated NCOR2-013 overexpression THP-1 macrophages. (F-G) The NCOR2-013 overexpression THP-1 macrophages and control cells were stimulated with LPS (500ng/ml) for 24h, and were infected or non-infected with *T. marneffei* spores (MOI = 10) for 24h and 48h. (F) qRT-PCR assays for TNF-ɑ and IL-1β expression in LPS-stimulated NCOR2-013 overexpression THP-1 macrophages. (G) *T. marneffei* CFU by microdilution spot assay to evaluate antifungal activity of LPS-stimulated NCOR2-013 overexpression THP-1 macrophages. All data are from three independent experiments, and show mean ± SD. An two-tailed Student’s t-test was used to determine significance, denoted by * (P < 0.05), ** (P < 0.01), and ns (not significant).

**Supplementary Appendix are provided online as separate Word documents. Table S1-2 lists human primer pairs used for RT-qPCR analysis. Table S3 shows the ChIP-qPCR primers sequences for the study**.

## Supplementary Appendix

### 1. RT-qPCR primers sequences for the study

**Table 1.**
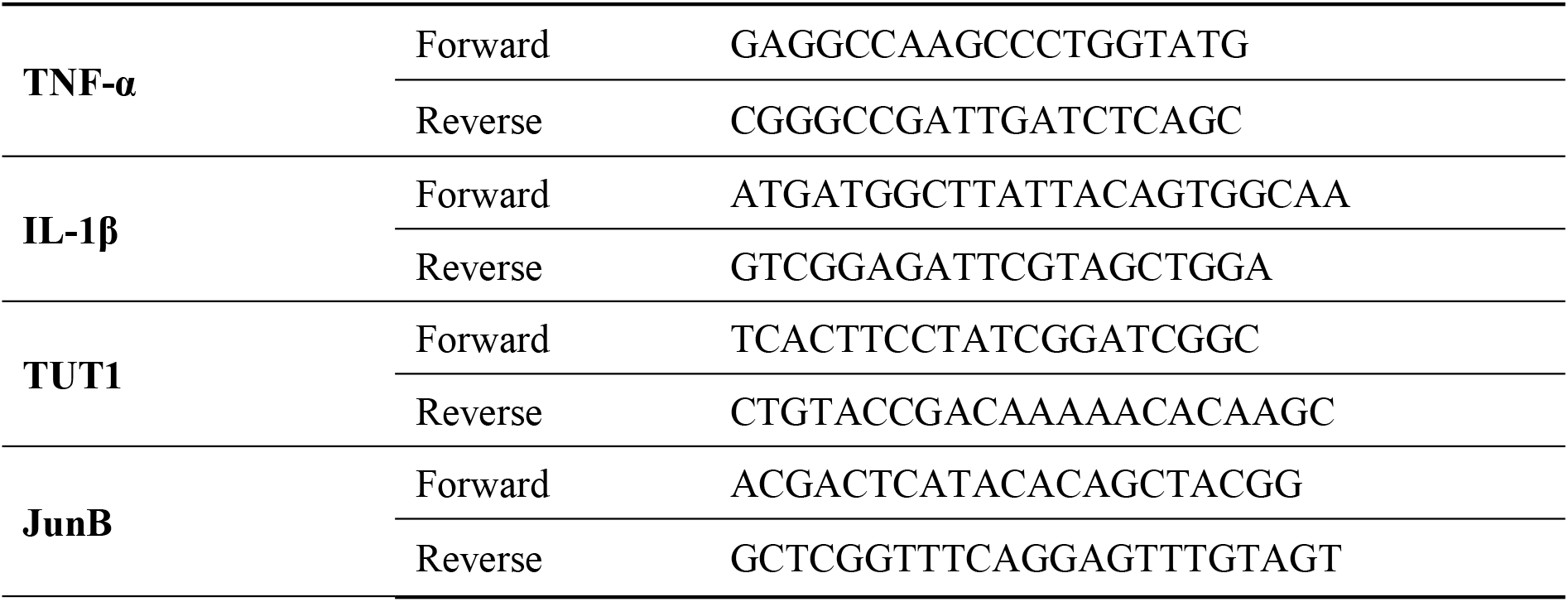
RT-qPCR primers sequences

### 2. Primer design principles for calculating NOCR2-013 expression levels

Exons 10 (ENSE00000997027) and 11 (ENSE00000997044) are all included among transcripts of NCOR2, which contain exon 12 (ENSE00000997028), while NCOR2-013 transcript contains only exons 10 and 11, but not exon 12. Therefore, two pairs of primers were designed based on the combined sequence of exons 10 and 11 and the sequence of exon 12. The expression levels of these two pairs of primers were detected by qPCR. Finally, the expression level of exon 12 was subtracted from the expression level of exon 10/11 to indirectly calculate the expression level of NCOR2-013 transcript.

**Figure.**
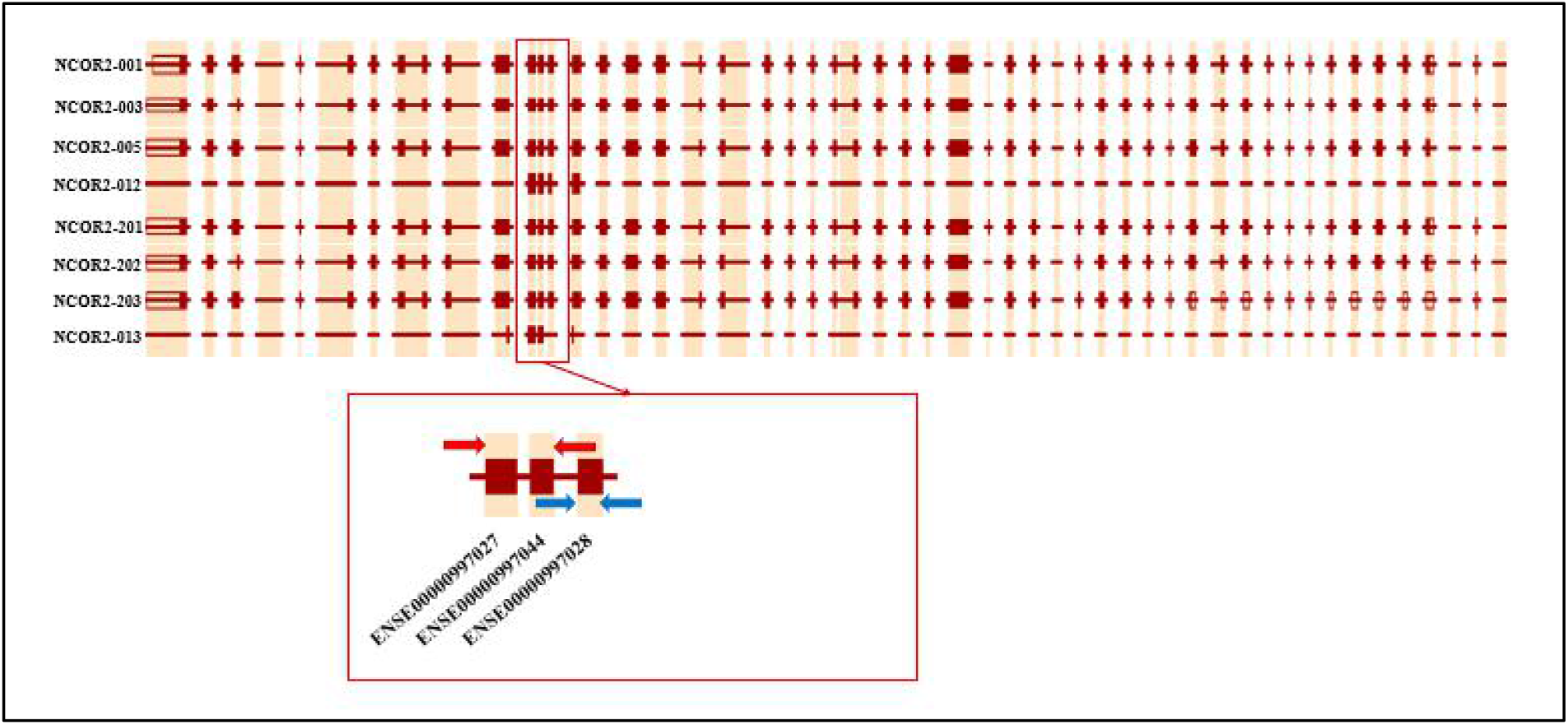
Primer design for NOCR2-013 expression levels.

**Table 2.**
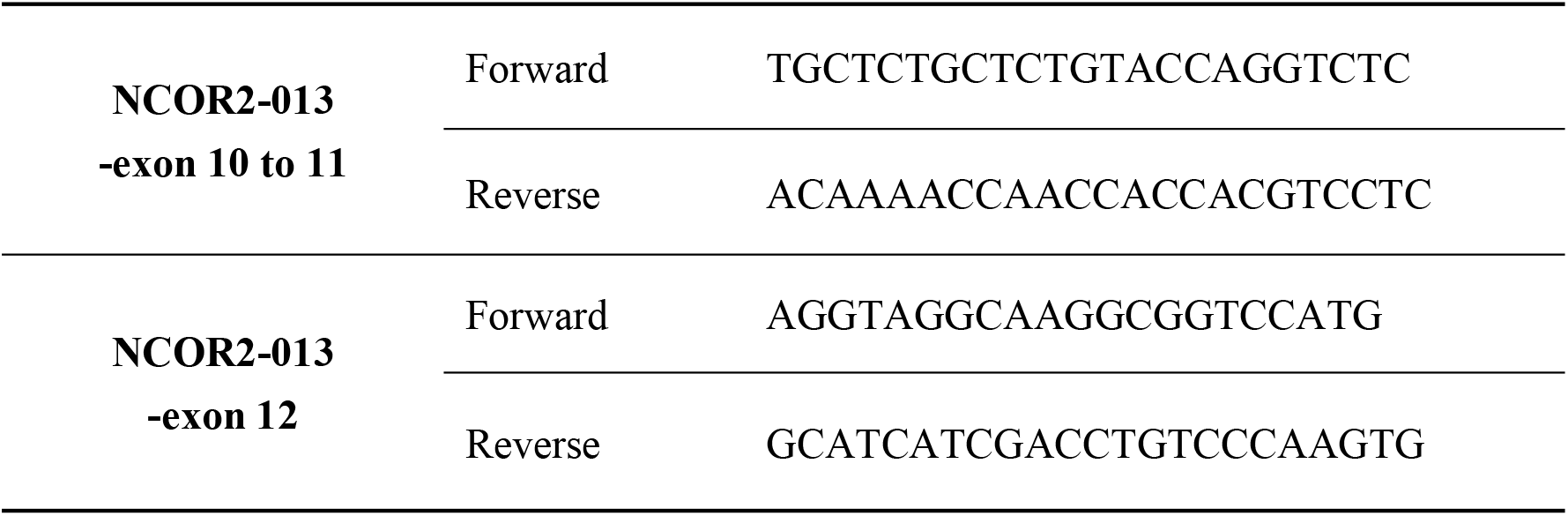
RT-qPCR primers sequences for NCOR2-013

### 3. ChIP-qPCR primers sequences for the study

**Table 3.**
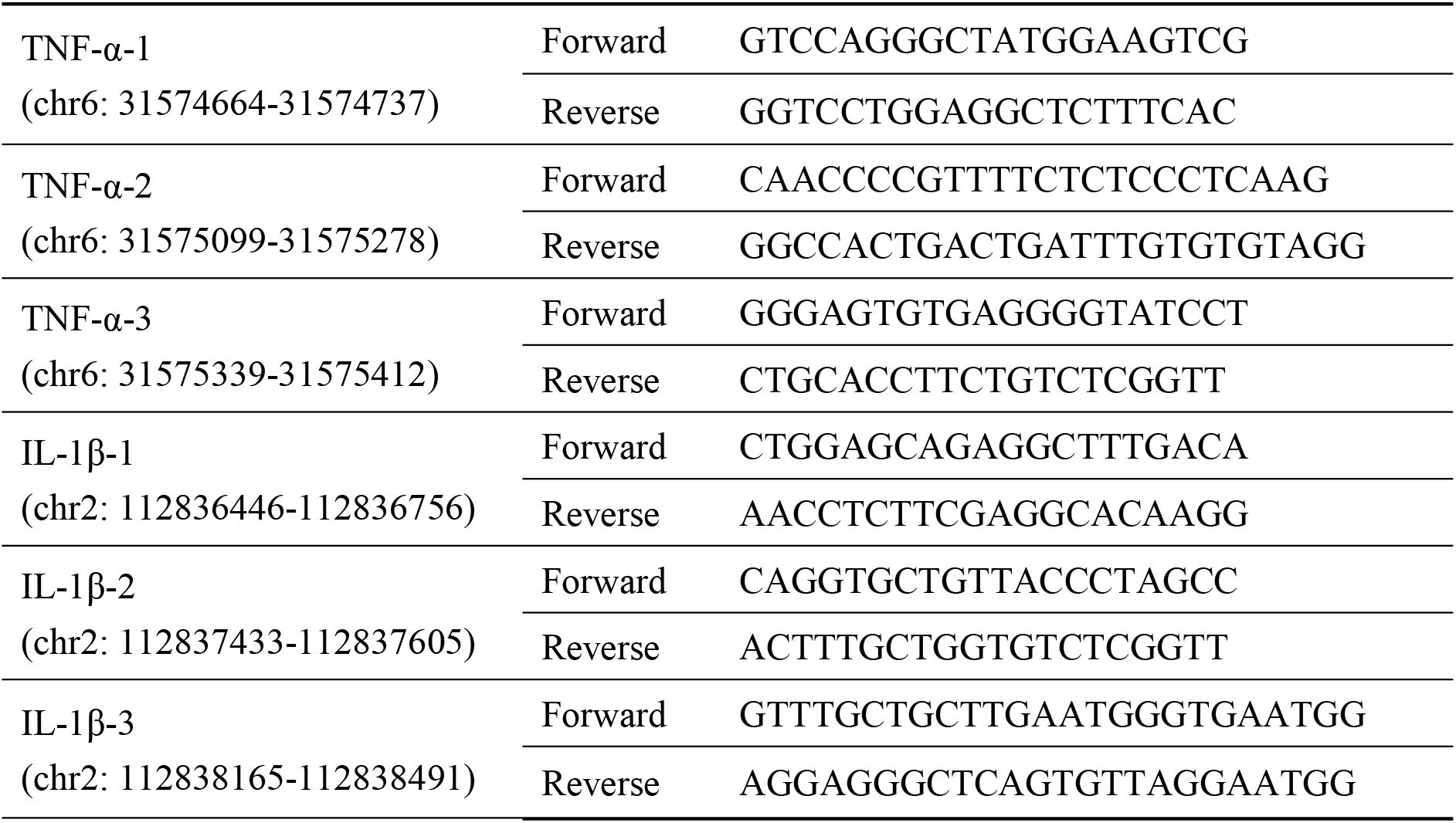
RT-qPCR primers sequences for promoter region of TNF-α and IL-1β

## Notes

### Competing Interest Statement

The authors have declared no competing interest.

